# Combining computational modeling and experimental library screening to affinity-mature VEEV-neutralizing antibody F5

**DOI:** 10.1101/2024.07.08.602599

**Authors:** Christopher A. Sumner, Jennifer L. Schwedler, Katherine Maia McCoy, Jack Holland, Valery Duva, Daniel Gelperin, Valeria Busygina, Maxwell A. Stefan, Daniella V. Martinez, Miranda A. Juarros, Ashlee M. Phillips, Dina R. Weilhammer, Gevorg Grigoryan, Michael S. Kent, Brooke N. Harmon

## Abstract

Engineered monoclonal antibodies (mAbs) have proven to be highly effective therapeutics in recent viral outbreaks due to their specificity and ability to provide immediate protection, regardless of immune status. However, despite technical advancements in the field, an ability to rapidly adapt or increase antibody affinity and by extension, therapeutic efficacy, has yet to be fully realized. We endeavored to stand-up such a pipeline using molecular modeling combined with experimental library screening to increase the affinity of a given antibody, F5, to recombinant E1E2 antigen from Venezuelan Equine Encephalitis Virus (VEEV) subtype IAB (TC-83). F5 is a monoclonal antibody with potent neutralizing activity against VEEV that was isolated from human bone marrow donors. F5 is known to bind to spikes on the surface of VEEV made up of a trimer of heterodimers of the glycoproteins E1 and E2. In this work we modeled the interaction of F5 with the E1E2 trimer of VEEV (TC-83) and generated predictions for mutations to improve binding using a Rosetta-based approach and dTERMen, an informatics approach. Modeling the structure of the complex was complicated by the fact that a high-resolution structure of F5 is not available and the H3 loop of F5 exceeds the length for which current modeling approaches can determine a unique structure. To overcome these challenges nine F5 structures with varying H3 loop conformations were generated using RosettaAntibody, PIGS (Prediction of ImmunoGlobulin Structure), and SWISS-Model and these base antibody structures were evaluated in docking trials to recombinant VEEV E1E2 based on relative binding affinity for several subtypes. The structure that gave the best agreement with the experimental trend in relative binding affinity was used for mutation analysis. A subset of the predicted mutations from both methods were incorporated into a phage display library of scFvs (single-chain variable fragments) and screened for binding affinity to the recombinant E1E2 antigen. Results from this screen were used to identify favorable mutations which were then incorporated into twelve human-IgG1 variants. All twelve variants showed increased binding relative to the parental F5 human-IgG1. The best case showed > 60x increased binding to recombinant E1E2 relative to the parental antibody, notably showing a drastic improvement of the Kd or “off rate” compared to the parental F5 IgG. These results demonstrate the ability of our methods to rapidly increase affinity and could be leveraged for increasing Ab binding breadth to additional viral variants.

## I. INTRODUCTION

In an increasingly interconnected world, the global risk of exposure to emerging pathogens makes the need for safe, potent therapeutic antibodies (Abs) ever more important for ensuring world health safety and security.[1, 2] The adaptive immune system can be trained to naturally develop affinity matured Abs for a known pathogen through vaccination or natural exposure, but over the past 40 years, the field of Ab research has expanded and deepened our understanding of Ab structure, function and engagement with the immune system, leading to our capacity to design and engineer novel, affinity-matured, therapeutics against emerging infectious agents. Optimizing binding affinity can improve potency and specificity, thereby reducing off target binding and side effects. Furthermore, developing highly potent, effective Abs can lower the effective dose and reduce the production costs. The engineering process of affinity maturation poses an interesting challenge as it involves a large sequence space of proteins 20^n^, where n is the number of amino acids in a protein. Random mutagenesis of scFv-based phage libraries typically results in 10^8^ – 10^10^ variants, and only a handful of mutations at critical complementarity-determining region (CDR) positions typically impact antibody affinity. Thus, random mutagenesis for affinity maturation is a slow and inefficient process [3, 4].

In-silico modeling holds promise to aid the process of affinity maturation through various Structure-Based Computational Design (SBCD) methods. Access to a large and ever-increasing protein database has facilitated the ability of computation to provide critical knowledge about fundamental interactions and the patterns of interacting amino acids in known structural motifs. We demonstrate with our modeling approach that computation can effectively screen mutants at a significantly higher rate than experimental approaches, thereby facilitating the requisite rapidity with which we must respond to new and emerging pathogens.

There are a variety of methods that may be utilized for SBCD of Ab-antigen (Ag) interactions [5]. Generally, any *de novo* method must involve prediction of the individual structures, prediction or design of their binding conformation(s), and prediction of binding strength for variations in Ab or Ag sequence for a given conformation or combination of conformations. RosettaAntibodyDesign [6] and OptCDR [7] both sample sub-structures of antibodies according to known canonical classes and graft them together. They then simulate mutating Ab residues followed by structural refinement with the mutations. Sequences are chosen with the best energies according to Rosetta energy and CHARMM energy respectively. OptMAVEn [8] simulates Variable, Diversity, and Joining gene (VDJ) recombination; the process by which lymphocytes randomly rearrange these gene segments to produce a broad repertoire of B cell receptors – Abs – and T cell receptors from a database of antibody structures including heavy chain (HC), and light chains (LC, kappa and lambda), then generates a docking ensemble and selects the structure with the best CHARMM energy. AbDesign aligns Abs and segments them according to points of maximum structural conservation, then grafts them together to generate many new backbones, followed by Rosetta sequence design [9]. Abs that already bind antigens, with known or predicted binding conformations, may also be redesigned to improve binding. For example, a given Ab variable region may be redesigned using a physics-based energy function [10], or Molecular Dynamics simulations combined with Monte Carlo sampling of different residues [11] to target the same or a different epitope within an antigenic domain to achieve increased affinity and therapeutic potency. In this work we demonstrate our process of redesigning the anti-VEEV Ab, F5, that is known to bind VEEV E1E2, and to improve upon its binding affinity and by extension efficacy.

Venezuelan Equine Encephalitis Virus (VEEV) is a New World alphavirus that can cause highly pathogenic neurological disease in humans and equines. The New World alphaviruses are considered potential biological weapons and are identified as Category B pathogens by the National Institute of Allergy and Infectious Diseases due to past weaponization, ease of producing large quantities of virus, and their highly infectious nature through aerosol exposure. There are currently no FDA-approved vaccines or drugs to prevent or treat neurotropic infections caused by VEEV and similar encephalitic viruses, [12–15], and thus we have an important opportunity to address these deficiencies by improving our ability to design and re-design Ab therapeutics to meet the challenge of these and future neurovirulent agents.

F5 is a monoclonal Ab that is potently neutralizing for VEEV and binds to domain A of the VEEV E2 envelope glycoprotein [16]. F5 was isolated using a phage display antibody library created from human bone marrow donors known to have circulating Abs for VEEV. We and others have shown that F5 provides 90-100% protection when given pre-exposure (-24 hours) and 70-90% protection, when dosed +24 to +48 hours post infection (hpi),in a lethal VEEV-TC83 model, with some variation depending on the route of infection whether aerosol, intranasal or subcutaneous (AE, IN or SQ). [16, 17]. Similarly F5 protects with 90-100% efficacy when dosed prophylactically (-24h) by SQ injection, AE, or IN administration, but that efficacy drops precipitously to 30%-40% at +24hpi, for SQ and IN exposure, and to 0% at +48hpi by IN route with fully virulent VEEV-TrD [16]. Interestingly, mice infected by aerosol challenge with VEEV-TrD and treated with F5 at +24hpi developed central nervous system infections but little or no clinical signs of disease and showed an 80% survival rate [12]. Accordingly, while F5 may be a strong candidate for prophylactic treatment of VEEV infection, its efficacy as a therapeutic at +24 and +48hpi is lacking. Thus, it was our goal to optimize F5 using our computational and experimental affinity-maturation pipeline to increase binding, and by extension enhance our therapeutic arsenal against encephalitic alphaviruses like VEEV.

In this work, computational modeling was combined with experimental library screening to improve the binding of the VEEV-neutralizing antibody F5 to recombinant E1E2 spike protein trimer of VEEV-TC83, an experimental live-attenuated VEEV vaccine derivative of virulent VEEV IAB subtype Trinidad donkey (VEEV-TrD). The modeling was particularly challenging because a high-resolution crystal structure was not available for F5, and the H3 loop of F5 is quite large (20 amino acids). Whereas the other CDR loops can be modelled from canonical structures, that is not the case for H3 due to its length [18, 19]. This challenge was addressed using published experimental data for the binding affinity of F5 to VEEV subtypes TC-83, IAB, IV, and V [16]. Nine F5 structures were docked to these various VEEV subtypes, and the complex that was most consistent with the published binding affinity data was selected for mutational analysis. Two modeling approaches were used to predict mutations to improve binding. The first approach was Sequence Tolerance [20, 21] within the Rosetta Protein Modeling Suite, a Monte Carlo-based tool for modeling structures and interactions of proteins. The second approach was dTERMen, an informatics approach [22]. Subsets of the predictions from both approaches (29 mutations at 13 sites) were screened using a phage library displaying scFvs that incorporated nearly all combinations of the predicted mutations. Here, we report the modeling procedures and the results of the library screening, which led to the identification of Abs that displayed increased binding to the E1E2 heterodimer by more than 60-fold, with significant improvement in the Kd (off rate) over the parental F5. Top candidate Abs displaying enhanced binding to the E1E2 heterodimer were tested for efficacy at binding and neutralization of virus *in vitro* as well as protection against lethal infection *in vivo*.

## II. COMPUTATIONAL METHODS

We modeled the interaction of F5 with VEEV-TC83 (subtype IAB) and F5 with subtypes IV, and V using the Rosetta Modelling Suite. Homology models for the surface E1E2 trimers of these viruses were generated from the VEEV-TC83 model 3J0C [23] along with cryoEM data for F5 bound to VEEV-TC83 (EMD-2645) [24]. The H3 loop of F5, comprising 20 residues, exceeds the length for which current modeling approaches can determine a unique structure [18, 25]. Nine structures were generated using RosettaAntibody, PIGS (Prediction of ImmunoGlobulin Structure), and SWISS-Model. Eight of the nine structures contained a kink in the H3 loop, since > 80% of known structures of CDR H3 loops contain a C-terminal kink [18]. These base antibody structures were docked against antigens of subtypes VEEV-TC83, IAB, IV, and V using Rosetta protocols Docking2 and Snugdock on the ROSIE server [26]. While a high-resolution structure for F5/VEEV is not available, the published cryoEM structure indicates that F5 binds near the center of the three E2 monomers in the E1E2 trimer [24]. Therefore, in the docking trials F5 was initially placed roughly 5 Å above the center of the trimer and models in which F5 bound to the trimer at locations away from the center were discarded. For each Ab/Ag pair one round of rigid docking (Docking2) was followed by 2 rounds of flexible docking (Snugdock). For the latter, the fast protocol was performed to avoid extensive remodeling of the H3 loop.

For predicting mutations to improve binding Rosetta-based approaches and an informatics approach were employed. For the former, both Sequence Tolerance [20, 21] and FlexddG [27] developed by Kortemme lab were used. Sequence Tolerance is an algorithm to predict the set of tolerated sequences for proteins and protein interfaces. From a single protein structure, a conformational ensemble is generated using Monte Carlo simulations involving backbone “backrub” and side chain moves. For each structure in the ensemble non-native sequences are generated by incorporating each of the twenty amino acids at sites defined by the user. The nonnative sequences are evaluated for both interface binding and fold stability and compared to the native sequence, and sequences with scores near to or better than the native sequence are saved as being “tolerated”. This is performed for the entire conformational ensemble and the frequency with which each amino acid is included within the set of tolerated sequences is reported. Sequence Tolerance was performed using the ROSIE public server.

FlexddG is a method for modeling changes in interfacial binding free energies upon mutation within the Rosetta Protein Modelling Suite [27]. It applies Rosetta’s interface ΔΔG module, ddg-monomer, to an ensemble of all-atom configurations to account for conformational flexibility. The configurational ensemble is generated using Rosetta’s “backrub” protocol that samples local side chain and backbone fluctuations for both partners of the protein-protein interface. In each instance, FlexddG evaluates ΔΔG for a single inserted mutation at a single specified site. CDR lengths were held constant and FlexddG was used to determine ΔΔG values for all twenty amino acids at each site targeted for mutation using recommended parameters of 35 structures, 5000 minimization iterations, and 35,000 backrub trials. For scoring, the generalized additive model (GAM) was used as described in Barlow et al. [27] The scores that result from FlexddG are in Rosetta Energy Units (REUs) using the Rosetta Talaris energy function. The python script for FlexddG analysis was downloaded from https://github.com/Kortemme-Lab/flex_ddG_tutorial and was employed with the Rosetta Software Suite (2018.48.60516). The Rosetta Relax function was also used in the modeling procedure. FlexddG was performed using high performance computational resources at Sandia National Laboratories (SNL).

At the point in time at which the experimental library was designed and fabricated, we did not have the capability to perform FlexddG analysis and so the mutations selected for inclusion in the library were generated using Sequence Tolerance for F5 sites that were in contact with the antigen in the model structure. However, after the library screening we employed FlexddG analysis for the same sites to compare predictions between Sequence Tolerance and FlexddG.

dTERMen [22] was also used to redesign all residues on the F5 antibody that contacted E1E2 in the structure selected for redesign. Briefly, given a template structure for design, dTERMen systematically breaks it down into constituent tertiary structural motifs (TERMs), then searches for low-RMSD matches for each TERM from a representative database made from the Protein Data Bank (PDB) to gather an ensemble of similar structures. Amino-acid sequence statistics in the resulting ensembles are then used to infer a second-order model of sequence space compatible with the template structure (i.e., a pseudo-energy model comprising self-energies for each possible amino acid at each design position and pair energies for all possible amino-acid pairs in interacting positions). Following the computation of self and pairwise pseudo-energies for these positions (the pseudo-energy table), linear programming-based optimization was used to find the sequence S_min that minimized the total pseudo-energy. Positions at which this optimal sequence differed from the antibody’s original sequence were taken as the first set of mutations considered [22, 28].

In addition, to search for mutations that may be specific to the antigen at hand, additional mutations were generated by optimizing the total pseudo-energy under a gentle “specificity gap”. A specificity-gap captures the difference between the energy of an amino acid at a given design position in the context of the surrounding residues (in this case nearby antigen residues), and the energy of that amino acid in similar backbone structures but other sequence contexts. Thus, it can be used to bias mutations at each design position for the sequence context of the antigen vs other sequence contexts. It is computed in a pseudo-energy table, wherein a specificity gap for amino acid α at position *i* is:

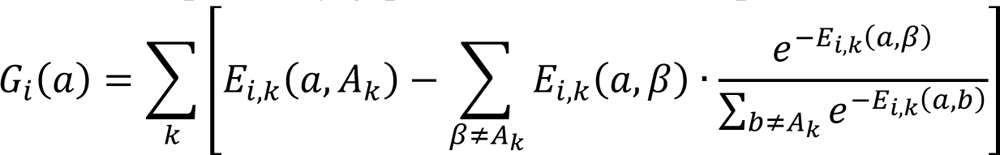

where *A_k_* is the amino acid at the antigen position *k* and *E_i,k_*(*α, x*) is the pair pseudo-energy between amino acid α at design position *i* and amino acid *x* at antigen position *k*. A second set of mutations were then designed as described above, except with the additional constraint that the total specificity gap of the designed position had to be below 8 units. This procedure resulted in a set of mutations that was larger than could be incorporated into the experimental library. To downselect a subset of mutations for inclusion in the experimental library from among the set of predicted mutations, input complexes were repacked with PyRosetta [29] using both sets of mutations, and then visually examined in PyMOL to manually screen for the most promising mutations based on shape complementarity and lack of clashing.

Subsets of mutations from the physics-based approach (Sequence Tolerance) and the informatics-based approach (dTERMen) were selected for inclusion in the experimental library.

## III. EXPERIMENTAL METHODS

### DNA Manipulation

pMTBip-TC83-E1E2 was constructed as follows and was adapted from other work to express recombinant E1-E2 heterodimer from Sindbis virus.[30] A double-stranded DNA fragment was commercially obtained (IDT) encoding the following TC83 structural protein coding sequences, E3 (1-59), E2 (1-344), and E1 (1-384). A Strep-tag II sequence (GGGSWSHPQFEKGGGG) was present in between the coding sequence for E2 and E1 in addition to a C-terminal hexa-histidine histidine tag. Trans-membrane regions of E2 and E1 were not included to produce soluble protein. This DNA fragment was assembled with agarose purified pMT/Bip/V5-HisA vector (Thermo), restriction digested with *BglII* and *XbaI*, and treated with phosphatase rSAP (NEB) using NEBuilder HiFi DNA master mix (NEB).

pSF-CMV-F5-hIgG1 and pSF-CMV-F5-λ were produced as follows. For the heavy and light chains of F5, double-stranded DNA gBlocks (IDT0 encoding the respective variable region was inserted upstream of the CH1-CH3 region of hIgG1 and the lambda CL1 . The genes were assembled with pSF-CMV vector restriction digested with *NcoI* and *BamHI* using NEBuilder HiFi DNA master mix (NEB).

### Protein Expression and Purification

VEEV-TC83 E1E2 glycoprotein was produced using a *Drosophila* S2 expression system. A Stable S2 cell line which expressed VEEV-TC83 E1E2 protein under an inducible promoter was generated as described in the technical literature provided by Gibco (Drosophila Schneider 2 (S2) Cells User Guide) . In brief, S2 cells were cultured in complete Schneider’s *Drosophila* Medium (10% heat-inactivated FBS, 1% penicillin streptomycin). S2 cells were transfected using a CaCl_2_ protocol. Cells were transfected at 3.0 x 10^6^ cells/mL with 19 μg of pMTBip-TC83-E1E2 and 1 μg pCoPuro in a 35 mm plate as described in the manufacturers protocol for 18 hours at 28°C. Cells were washed and media exchanged, then allowed to grow in the absence of selection agent for 48 hours. Cells were then expanded in the presence of 7 μg/mL puromycin. Stable polyclonal S2 cells were expanded and adapted to EX-CELL 420 serum-free media (Sigma) supplemented with 0.1% Pluronic F-68 over several passages with shaking. Protein expression was induced at a cell density reached 6 x 10^6^ cells/mL with 500 μM CuSO_4_.

Protein was allowed to express for 5 days after which the supernatant containing recombinant E1-E2 protein was harvested. Cells were centrifuged at 4,000 x g for 30 minutes at 4°C. Supernatant was then clarified by 0.45 μm filter prior to application to a 5 mL HisTrap Excel column (Cytiva) equilibrated with 20 mM Tris (pH 8.0), 300 mM NaCl. The column was washed extensively with buffer followed by washing with equilibration buffer with 40 mM imidazole. E1-E2 complex was eluted from the column with equilibration buffer supplemented with 500 mM imidazole. Fractions containing protein were dialyzed against 20 mM CHES (pH 9.5), 200 mM NaCl overnight at 4°C. Protein was concentrated and applied to Superdex 200 Increase 10/300 GL (Cytiva) equilibrated with 20 mM CHES (pH 9.5), 200 mM NaCl. Fractions containing E1-E2 complex were concentrated, aliquoted, and snap frozen with liquid N_2_. Protein was stored at -80°C.

### Library fabrication

To incorporate desired mutations at specific sites in the CDRs, codons were chosen using http://guinevere.otago.ac.nz/cgi-bin/aef/AA-Calculator.pl. The oligos were designed to include as many of the prioritized predicted mutations as possible within a total diversity of ∼ 10^8^, while avoiding stop codons and minimizing off-target residues. 20-25 nt homology overlap on both 3’ and 5’ ends was added for efficient annealing.

For generation and validation of the parental F5 scFv phagemid, the parental F5 scFv sequence was cloned into pAP-III_6_ vector [31] in VH-VL orientation with (GGGS)_4_ linker. This cloning event fused F5 scFv to the coat protein III of bacteriophage M13. The phagemid was transformed into NEB ® Turbo chemically competent *E. coli* strain and phage supernatants were prepared as described after cell growth and transduction with M13-K07 helper phage (NEB) at multiplicity of infection (MOI) of 10. Binding of F5 phage supernatants to VEEV E1E2 heterodimer was determined by ELISA.

Randomly mutagenized scFv library was produced as described in .[32]. Targeted mutagenized scFv library was produced using modified Kunkel mutagenesis method as described in .[33]. Briefly, to produce ssDNA phagemid template for library generation, Eco29K I restriction sequences (CCGCGG) were introduced in each CDR of the F5 antibody sequence. The ssDNA template was produced in CJ236 strain (NEB) as previously described [32]. Oligo mixture comprising a mix of WT and mutagenized sequences for each CDR was annealed to the ssDNA template, followed by the extension and ligation reactions to generate dsDNA products. The newly generated mutagenized plasmids were transformed into AXM688 *E. coli* strain [17] expressing Eco29KI restriction nuclease for removal of all non-recombinant products from the library. To validate the presence and proportion of each designed mutation, 48 single colonies were selected and F5 scFv variants were obtained by colony PCR and analyzed by Sanger sequencing.

To initiate the first round of screening, ELISA plates (Nunc Maxisorp) were coated with purified E1E2 VEEV-TC83 heterodimer(100 µl, 10 µg/ml) overnight at 4°C. Phage library was added to the plates at approximately 1^1^x 10^11^ transforming units and incubated for 2 hours at RT. The plates were then extensively washed and bound phage eluted and propagated in *E.* coli to be used for the next round of screening. The screen proceeded essentially as described [34] through 3 rounds of selection with decreasing concentrations of the E1E2 heterodimer (3 µg/ml in round 2 and 1 µg/ml in round 3). After round 3, single colony phage was prepared, and phage ELISA performed as described [34] against E1E2 heterodomer coated on ELISAELISA plates at 3 µg/ml (100 µl/well). Mutational analysis of the selected F5 variants was performed by PCR from the glycerol stock of single phage colonies followed by Sanger sequencing.

### Virus Stocks

Vero cells (African green monkey kidney, ATCC CCL-81) were cultured in minimum essential medium alpha (alpha MEM) supplemented with 10% fetal bovine serum (FBS), 100 units/ml penicillin, and 100 μg/ml streptomycin (Invitrogen, 15070063), at 37 °C in 5% CO2. VEEV--TC83 was obtained from the NIH Biodefense and Emerging Infections Research Resources Repository, NIAID, NIH (NR-63) and amplified in supplemented alpha MEM. TC83 stocks were amplified in Vero cells infected at a MOI of 0.1, 40 hours post infection virus was harvested and cellular debris removed by centrifugation 1000 x g for 5 minutes. Clarified supernatant containing TC83 was either snap frozen as a crude preparation with liquid nitrogen and stored at -80°C or further purified by density ultracentrifugation. In brief, crude preparation of virus was added on top of a cushion of 20% sucrose in PBS and centrifuged in a SW 40Ti rotor (Beckman) at 30,000 rpm for 1.5 hours. Media and sucrose were then carefully aspirated and purified virus was reconstituted in PBS and stored at -80°C. Viral titers were determined by plaque assay.

### ELISA

Binding of IgG candidates from display screening was evaluated against both recombinant E1-E2 glycoprotein and live VEEV TC83 by immunosorbent assay. For recombinant E1-E2, glycoprotein was diluted in 100 mM NaCO_2_ (pH 8.3) 150 mM NaCl and immobilized on 384 well immulon HBX plates (Thermo) overnight at 4°C. For ELISA with live virus, TC83 was diluted to 10^7^ PFU/mL in PBS and 25 μL added to each well for overnight incubation at 4°C. After washing with PBST, plates were blocked for 2 hours at 25°C with Pierce (PBS) Protein Free blocking buffer. After removing block, serial dilutions of IgG candidates were prepared in blocking solution and allowed to incubate for at least 2 hours in the plate with gentle shaking. The plate was washed and 1:15,000 HRP-conjugated goal anti-human (Thermo) was added. Plates were developed with TMB Ultra (Thermo) and the reaction stopped by addition of equal volume 2 M H_2_SO_4_. Absorbance was read at 450 nm.

### Plaque Reduction Neutralization Test (PRNT) Assays

Neutralization assays to determine the plaque reduction neutralization for candidate IgG antibodies were performed as follows. Serial dilutions of each antibody in alpha MEM were mixed with equal volume of 25 PFU TC83 virus to form antibody-virus complexes. This mixture was preincubated at 37°C with 5% CO2 for 1 hour prior to addition of Vero E6 cells at ∼80% confluency in 12 well plates and allowed to incubate for 30 minutes. Overlays of 1.5 mL 0.5% agarose in MEM were added to each well and allowed to solidify. After 36-40 hours plates were rinsed with PBS and stained and fixed with 0.25% crystal violet (w/v) and 0.2% paraformaldehyde (v/v) for 30 minutes prior to extensive washing with water. Plaques were counted and EC50 values were determined using GraphPad software using the following equation: Y=Bottom + (Top-Bottom)/(1+(IC50/X)∧HillSlope).

### Biolayer Interferometry

Kinetic parameters for each anti-VEEV IgG antibody were determined using an Octet^®^ RED384 (Sartorius). Each IgG antibody was immobilized on anti-human Fc sensors in 10 mM phosphate (pH 7.4), 300 mM NaCl, 1 mg/mL BSA, 0.02% NP-40. VEEV TC83 E1E2 was used as the analyte and sensograms were fit to 1:1 global kinetics.

### Animal Studies

All animal work was approved by the Lawrence Livermore National Laboratory Institutional Animal Care and Use Committee. All animals were housed in an Association for Assessment and Accreditation of Laboratory Animal Care (AAALAC)-accredited facility. C3H/HeN female mice (Charles River) ranging in age from 4-8 weeks were inoculated intranasally with 5x10^7^ pfu VEEV-TC83 under anesthesia (4-5% isoflurane in oxygen). Animals were dosed with therapeutic antibodies at +24 or +48 hours post infection via intraperitoneal injection (4mg/kg in PBS). Mice were monitored daily for signs of morbidity up to 14 days post infection (dpi). Animals were humanely euthanized by CO_2_ asphyxiation upon signs of severe disease. Survival data were analyzed for significance using the Log-rank (Mantel-Cox) test using Prism software (GraphPad, La Jolla, CA).

## IV. RESULTS

### In-Silico Mutation Analysis

Since the cryoEM structure EMD-2645 indicates that F5 binds near the center of the three E2 monomers in the E1E2 trimer [24], and considering the large length of the H3 loop, it seems likely that the H3 loop interacts with and inserts into the open space in the center of the E1E2 trimer. Since a high-resolution structure for F5 was not available, nine F5 models with varying H3 conformations were generated using RosettaAntibody, PIGS, and Swiss Model (Figure S1). The H2 loop structure was very similar in 8 of the 9 models as shown in Figure S1a, whereas the structure of the H3 loop varied greatly among the nine models. To select among these models the nine structures were used in docking trials against IAB (TrD and TC-83), IV, and V antigens using Rosetta protocols Docking2 and Snugdock. The results, shown in Figure S2, were evaluated based on the trends in relative binding affinity for each of the subtypes. A prior study reported the relative binding as TC-83 ∼ TrD > IV > V [16]. The RosettaAntbody1 structure gave results that were most in line with the experimentally observed trend. For that structure the docking results gave the binding affinity as TC-83 ∼ TrD > IV. The docking results for binding to subtype V were not consistent with the reported experimental trend, as no detectable binding was reported at 100 ug/ml for subtype V in the experimental study. Nevertheless, since the results were consistent with the trend for the other subtypes this structure was selected and further refined in docking trials using the Relax function followed by two additional rounds of flexible docking using Snugdock on ROSIE (fast protocol). This final structure, shown in Figure 1, was used to generate predictions for mutations to improve binding.

**Figure 1.**
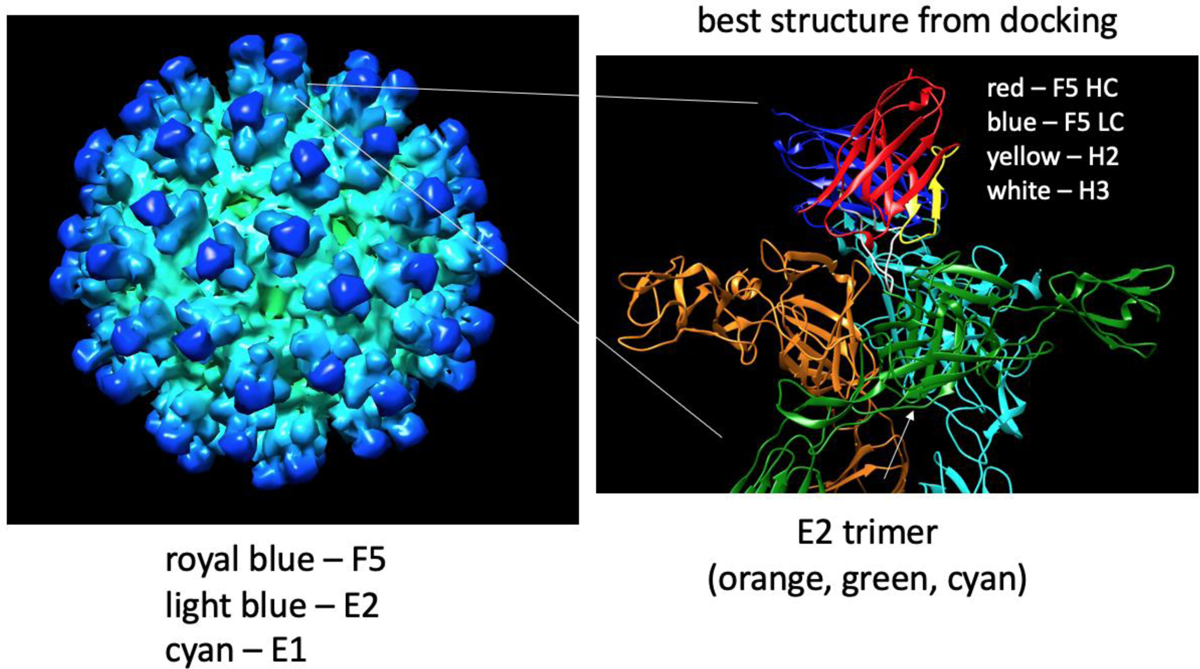
The structural model used for mutational analysis by Sequence Tolerance, FlexddG, and dTERMen.

The results of Sequence Tolerance Analysis for the final refined F5-VEEV structure are given in Figure S3. Considering the sites in direct contact with the antigen, or only 1 site removed from direct contact, the analysis indicated that at 24 sites mutations were preferred or tolerated at comparable frequency to the parent residue. Among these, 19 mutations at 13 sites were included in the experimental library, indicated in gold in Figure 2.

**Figure 2a.**
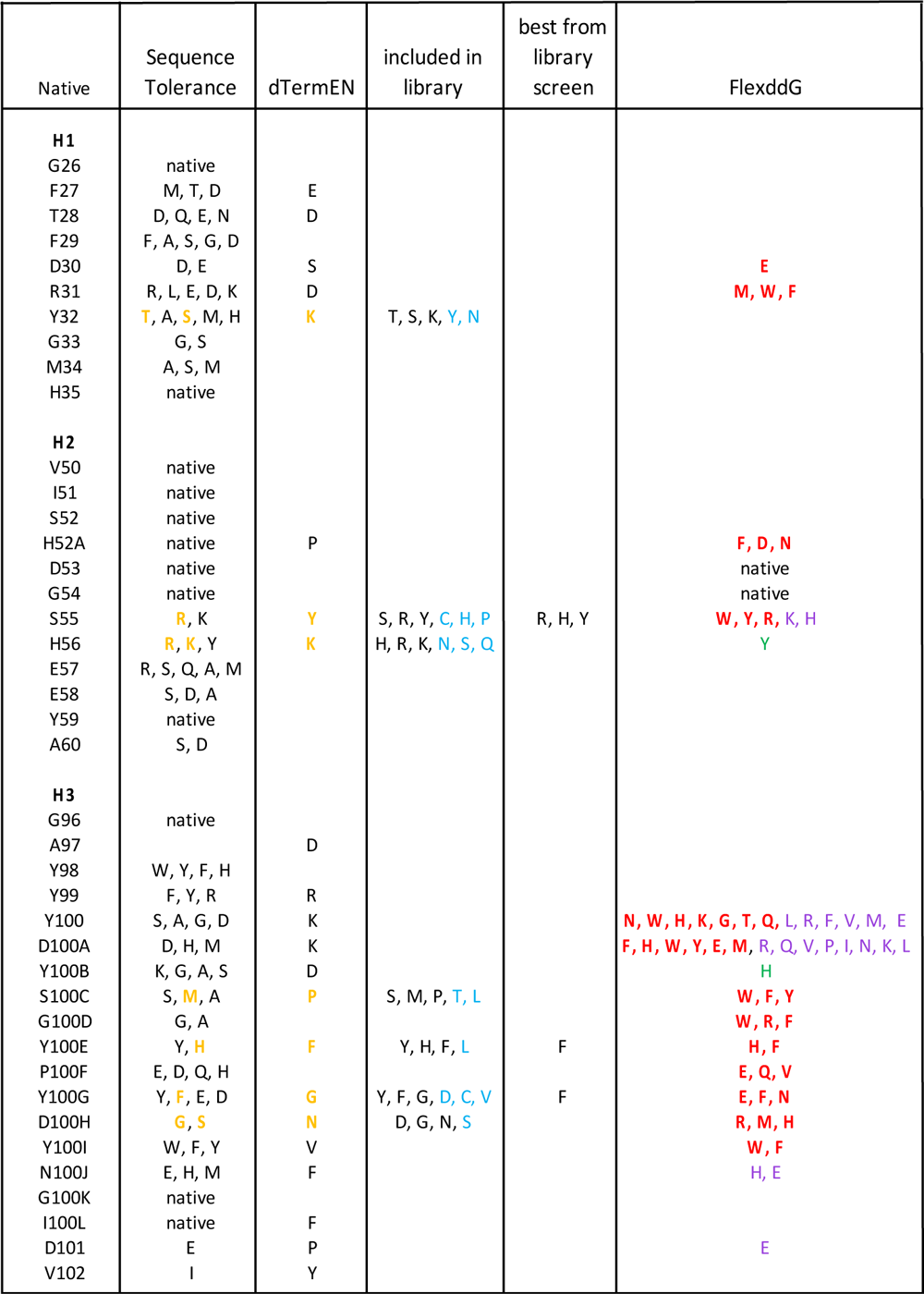
Heavy chain mutation predictions and results of library screening. The predicted mutations from Sequence Tolerance, dTERMen, and FlexddG mutations are shown along with mutations that were highly selected in the library screening. The mutations that were included in the library are shown in the fourth column. Mutations that were not predicted but were a consequence of including the predicted mutations are shown in cyan in the fourth column. The mutations predicted by Sequence Tolerance and dTERMen that were included in the library are shown in gold. For the FlexddG predictions, mutations predicted to be strongly favorable (> 1.0 REU), moderately favorable (0.5 REU to 1.0 REU) and marginally favorable (0.2 to 0.5 REU) are shown in red, purple, and green, respectively.

**Figure 2b.**
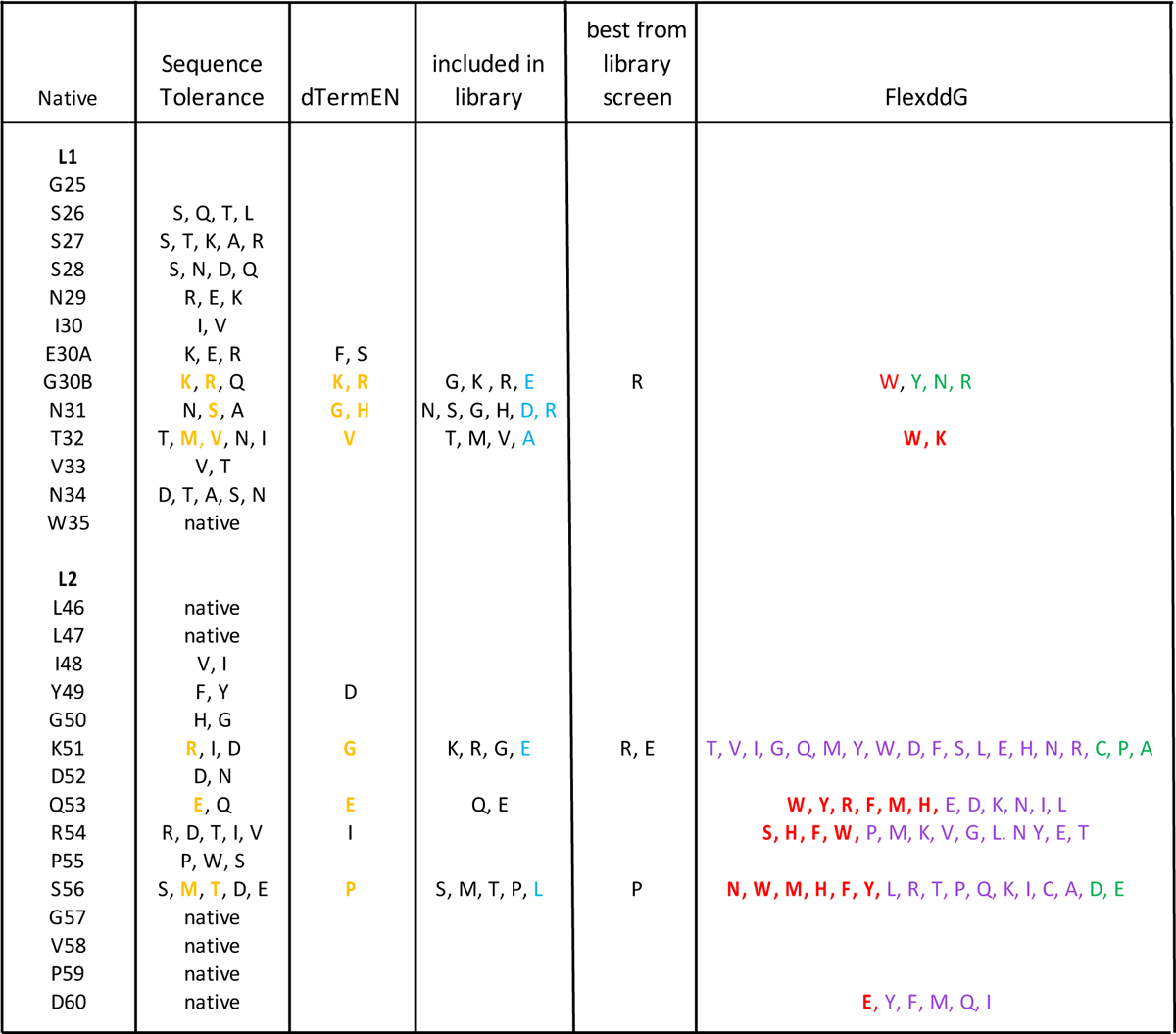
Light chain mutation predictions and results of library screening.

dTERMen was also used to predict favorable mutations in the final refined F5-VEEV structure. Two sets of predicted mutations, one based on pseudo-energies alone and another based on pseudo-energies with a light specificity cutoff (see Methods), are given in Figures S4 and S5. Considering the sites in direct contact with the antigen or only 1 site removed from direct contact, the results indicated favorable mutations at 25 sites. Among these, 15 mutations at 13 sites were included in the experimental library. Downselection was required due to constraints on the size of the experimental library. The mutations included in the experimental library are, highlighted in gold in Figure S6 and in Figure 2. Of the 15 mutations, 10 were unique to dTERMen and 5 were in common with the predictions from Sequence Tolerance.

After the experimental screening, FlexddG was performed for all sites at which mutations were included in the experimental library. The results are given in Figure 2, where bold red indicates favorable ΔΔG values greater than 1 Rosetta Energy Unit (REU), purple indicates favorable ΔΔG values between 0.5 and 1.0 REU, and green indicates favorable ΔΔG values between 0.2 and 0.5 REU.

### Library screening

Two phage libraries were tested for binding to the recombinant VEEV (TC-83) E1E2. Each library had a diversity of roughly 4 x 10^8^. One library was generated by randomly introducing mutations throughout the entire scFv. The second library was generated by introducing specific mutations from the in-silico modeling. Despite the large library size of 4 x 10^8^, not all predicted beneficial mutations could be screened. While larger libraries have been routinely made by many labs, including ours, we aimed here to generate a library with a smaller number of mutations that interrogates nearly all possible combinations of the mutations chosen for inclusion [32]. A total of 19 mutations from Rosetta and 15 mutations from dTERMen (10 unique and 5 in common with Rosetta) at 13 sites were included in the directed library. These mutations were chosen based on the anticipated impact on binding as well as considerations from codon usage such as avoiding stop codons and minimizing the number of off-target amino acids generated at each site of mutation. In addition to the desired mutations, an additional 21 mutations were present as a necessary consequence of the codons used to incorporate the desired mutations.

#### Screening Results - Directed library

After 3 rounds of affinity selection 68 clones were identified and sequenced. Based on the ELISA signal intensity shown in Figure 3, 29 unique clones were identified as having improved binding relative to the parental F5, 14 clones had comparable binding to the parental, and 23 clones had lower binding than the parental. Mutations were identified as beneficial, neutral, or detrimental based on their distribution among the clones in the three categories (improved, comparable, or lower binding relative to parental). The results are tabulated in Figure S7-S9. To assign a mutation as being clearly beneficial, we used the criteria that a mutation must have appeared in greater than three clones and roughly 50% or greater of all instances of that mutation must have occurred within the clones that gave higher binding than parental. We note that this latter criterion considers all mutations independently, even though other mutations in a clone will likely affect binding. By that criterion, nine mutations at six positions were identified as highly likely to be beneficial for improving binding ((Figure 4). Three of the nine were uniquely predicted by the Rosetta method, three were uniquely predicted by dTERMen, one was predicted by both methods, and two were off-target amino acids that were present because of the nucleotides required for the targeted amino acids. The latter demonstrates that there is value in simply identifying the interacting amino acids and focusing library diversity on those sites. Another important point is that simply combining all predicted mutations into a single construct did not lead to successful clones. For the 28 clones that had improved binding to the target, the number of mutations ranged from 2 to 6 with a mean of 3.8.

**Figure 3.**
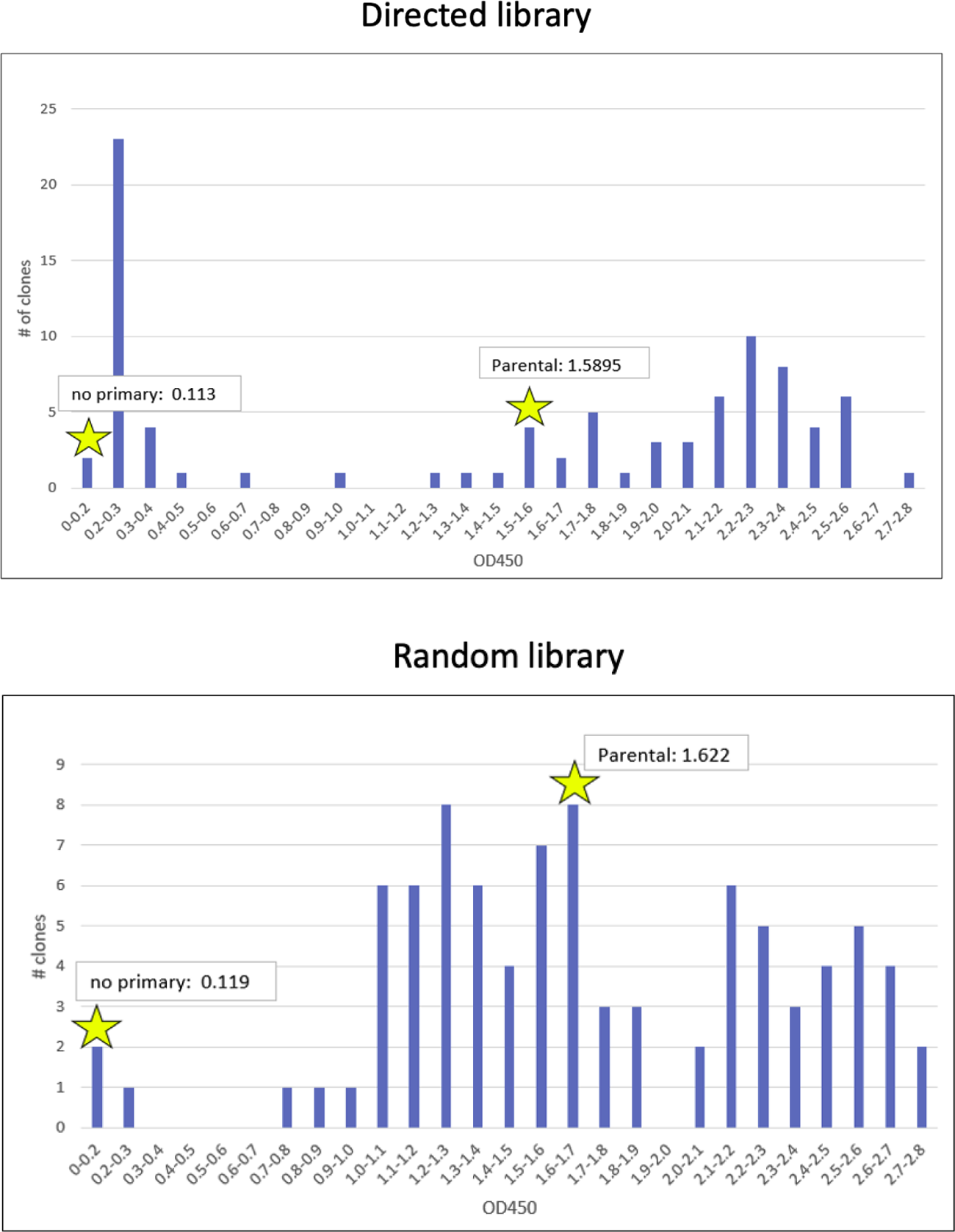
ELISA results for binding of clones to the E1E2 heterodimer for the two libraries.

**Figure 4.**
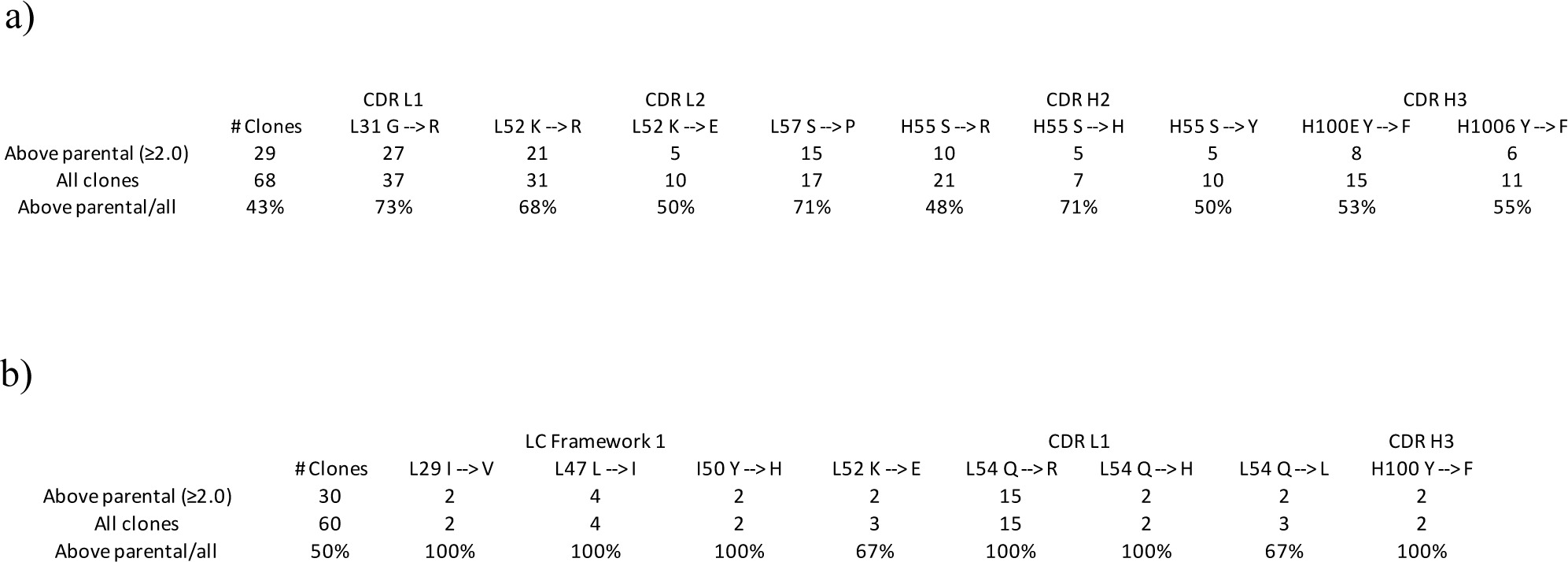
Summary of beneficial mutations for a) directed library, and b) random library.

Regarding FlexddG, four of the experimentally-determined favorable mutations were identified within the most strongly favored range (> 1 REU), four of the experimentally-determined favorable mutations were identified within the range 0.5 – 1.0 REU, and 1 of the experimentally-determined favorable mutations was identified within the most weakly favored range 0.2 – 0.5 REU.

#### Screening Results - Random library

Using the same criteria, two mutations were identified in the random library as improving binding. One of these occurred in the framework region and the second occurred in CDR L1. The latter was distinct from the beneficial mutations identified in the directed library (Figure 4).

### Binding studies reveal enhanced binding affinity for twelve new F5-derived Abs

Our top twelve antibody candidates were designed to contain multiple combinations of the beneficial mutations identified in our experimental library screens and are listed in Figure 5. These Abs were tested for binding to the recombinant TC-83 E1E2 heterodimer by ELISA and by bilayer interferometry (BLI). The ELISA results, shown in Figure 6, indicate increased binding for all twelve of our mutant Abs (SNL1-1 - SNL1-12) relative to the parental F5 antibody (SNL1-13). Of the twelve mutational combinations tested, some yielded only a very modest improvement on binding to TC-83 (SNL1-4 and SNL1-8). Our top binding candidates SNL1-1 and SNL1-2, which included the most potent beneficial mutations with the additional framework mutation included in SNL1-1 but not SNL1-2, showed a substantial improvement in binding relative to the parental F5 (SNL1-13).

**Figure 5.**
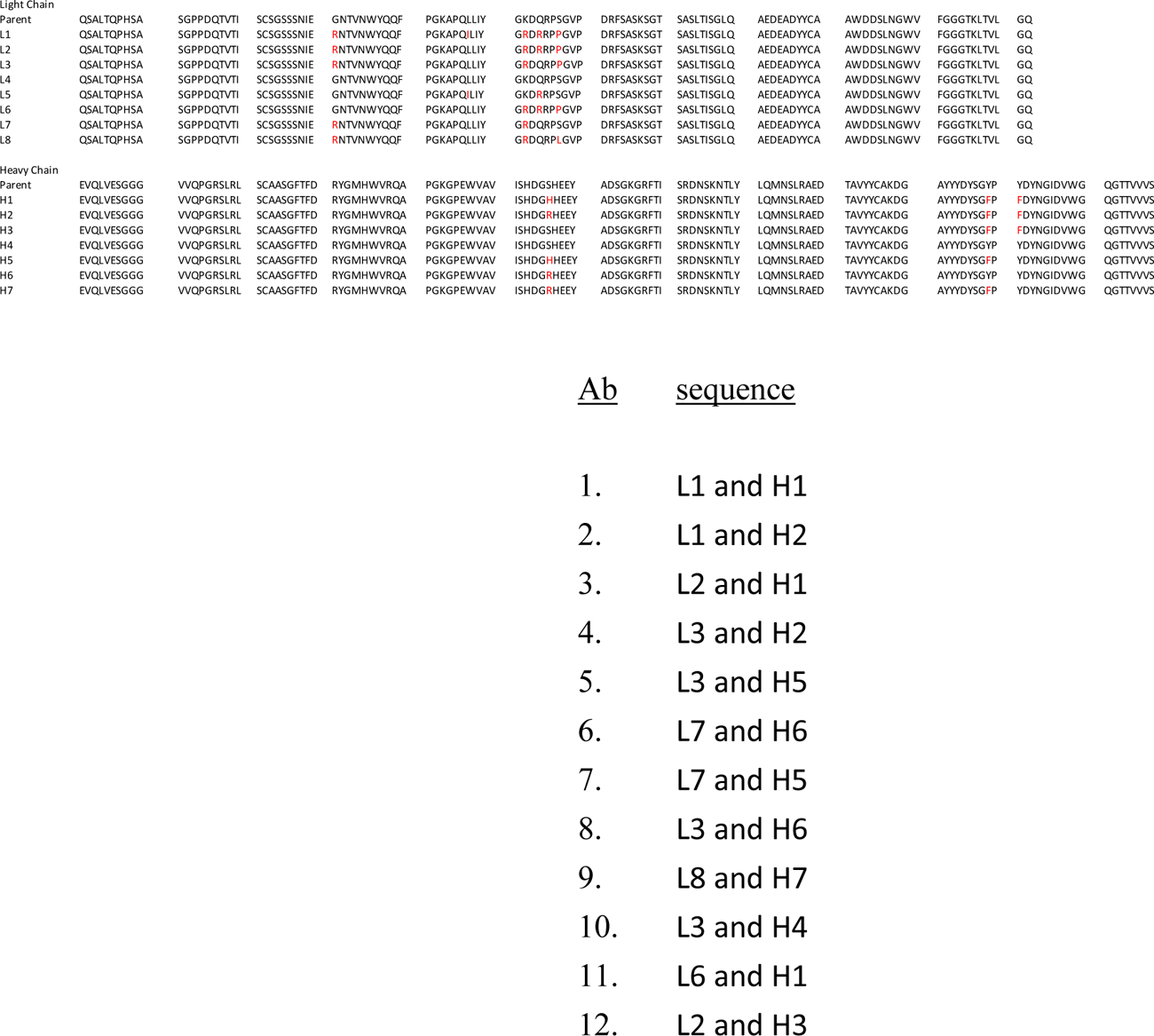
Sequences of twelve fabricated IgGs that incorporate various combinations of the mutations identified as favorable for binding.

**Figure 6.**
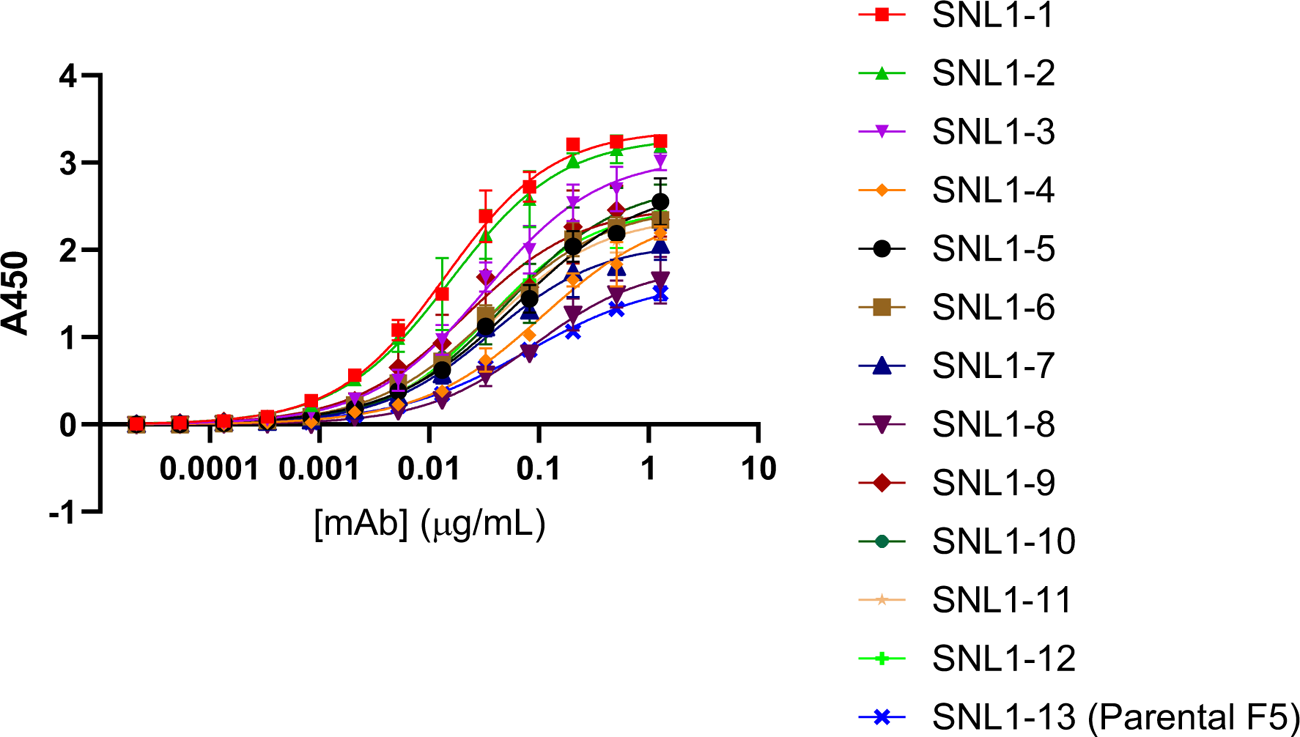
ELISA for IgGs binding to recombinant TC-83 E1E2 heterodimer.

BLI results for our designer Abs binding to recombinant E1E2 , shown in Figure 7, agree well with the ELISA data and provide quantitative values for binding affinity. Of the mutant candidates tested, the largest improvement in binding relative to the parental occurred for SNL1-1 and SNL1-2, showing a 63 fold and 3.9 fold decrease in KD relative to F5 (SNL-13) in our assay. When taking a closer look at the BLI binding data, we noticed that while there is some modest improvement in the Ka (association) for many of the mutants tested, all fell within a relatively close range (2.90x10^5^ -2.17x10^5^ M^-1^s^-1^), suggesting that the mutations made did not drastically affect the association rate of F5 with TC-83 E1E2. Additionally, the top candidates SNL1-1 and SNL1-2 were not at the extreme end of this range with values of 2.33x10^5^ and 2.44x10^5^ M^-1^s^-1^ respectively, suggesting that improvement in association was not the main driver of their selection. Interestingly, while all mutant candidates showed improvement in the Kd (dissociation) relative to the parental F5, most mutants show some modest improvement (≤4.8 fold decrease in Kd) while SNL1-1 showed a drastic improvement in Kd (60.3 fold decrease). Interestingly, this suggests that the combination of binding enhancement mutations as well as the framework mutation included in SNL1-1 result in a modest increase in association rate, but a significant improvement in dissociation rate, resulting in a more stable antigen-antibody interaction (Figure 7).

**Figure 7.**
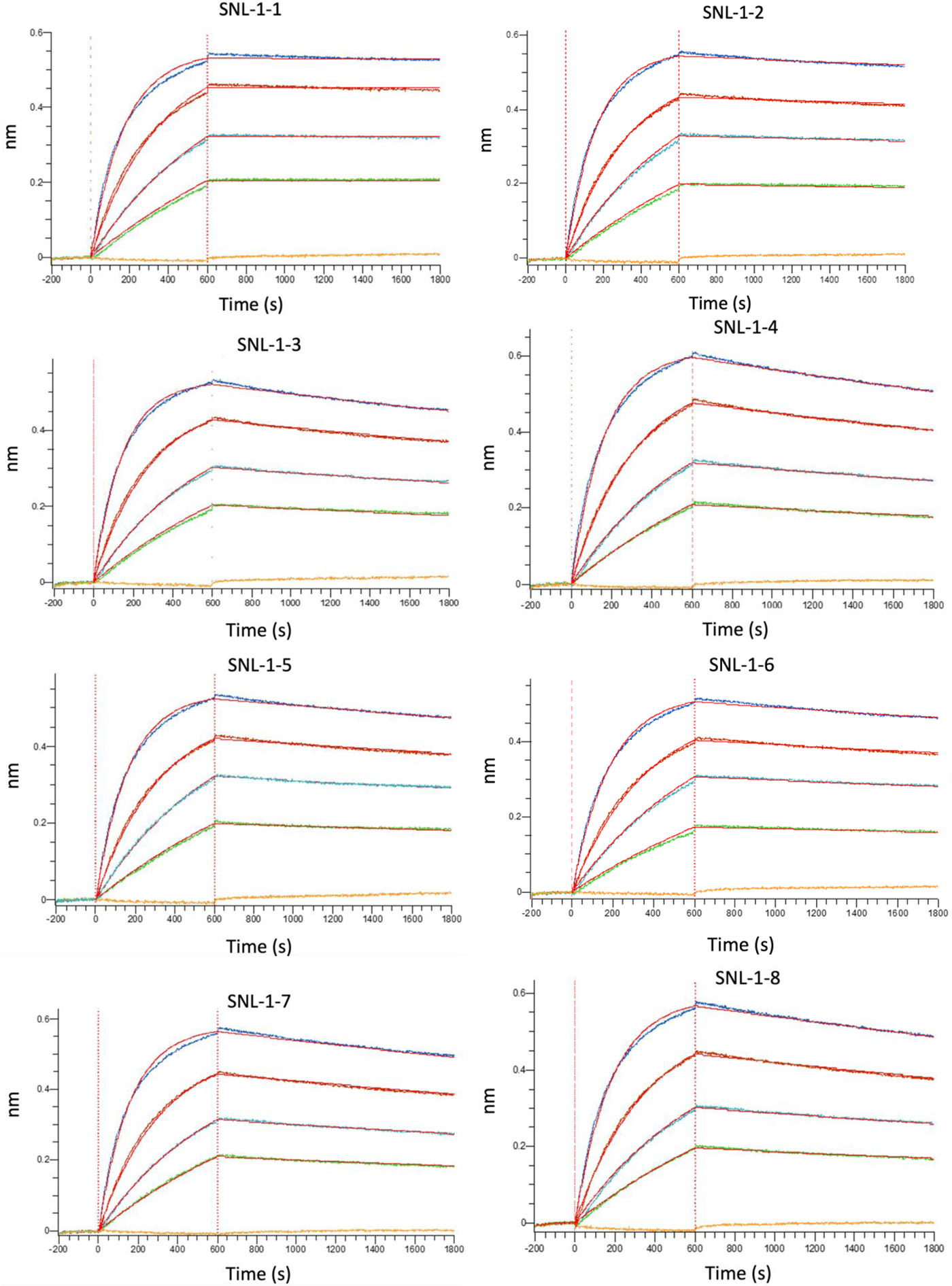
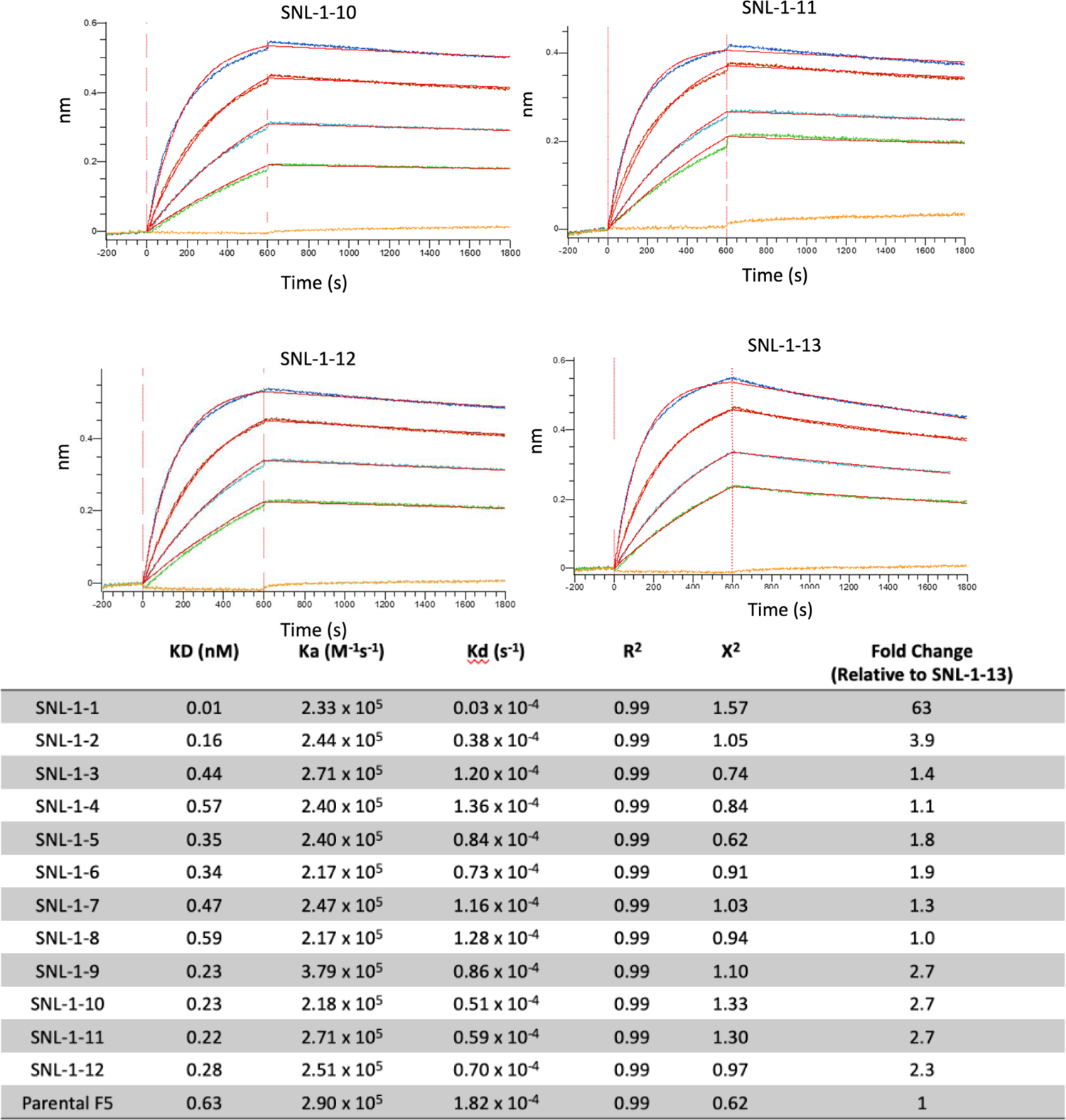
Bilayer interferometry for the IgGs binding to the TC-83 E1E2 heterodimer.

### Despite enhanced binding, SNL 1-1 does not improve efficacy against VEEV-TC83

To determine the impact of enhanced Ab-binding to recombinant E1E2 on our Abs capacity to bind live virus, we performed a plaque reduction neutralization test (PRNT). EC_50_ values are listed in Table 1. The raw data for percent neutralization versus antibody concentration are given in the Supporting Information (Figure S11). The results show no statistical improvement in neutralization for all twelve designed antibodies over the parental. Despite the lack of improved neutralization capacity, non-neutralizing Abs have shown efficacy against VEEV, so we tested the ability of our top candidate, SNL1-1, which displayed the highest binding affinity for E1E2, to protect against infection with VEEV-TC83 *in vivo* using an established mouse model of lethal infection (PMID 18313150). Since F5 has previously been reported to be up to 100% effective at preventing lethal infection when delivered prophylactically,[16] we assessed efficacy of SNL1-1 when delivered therapeutically at +24 and +48hpi. The results of the survival experiments are shown in Figure. Treatment with both SNL1-1 and the parental F5 antibody (SNL1-13) resulted in significant protection against infection when delivered at +24hpi (Figure 9A). Protection against infection was also seen at +48hpi (Figure 9B), although increased survival was not statistically significant at this timepoint. There was no statistical significance in survival observed between SNL1-1 and F5 treated groups at either time point.

**Figure 8.**
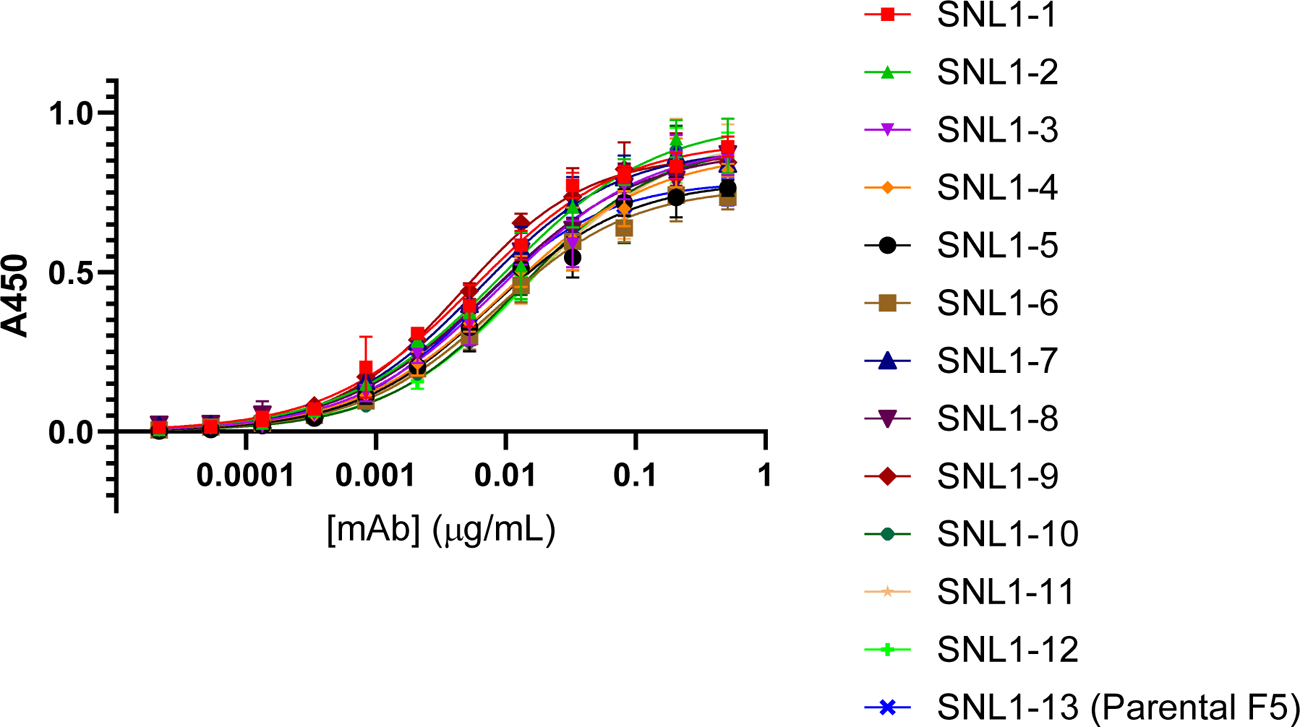
ELISA for IgGs binding to inactivated TC-83.

**Figure 9.**
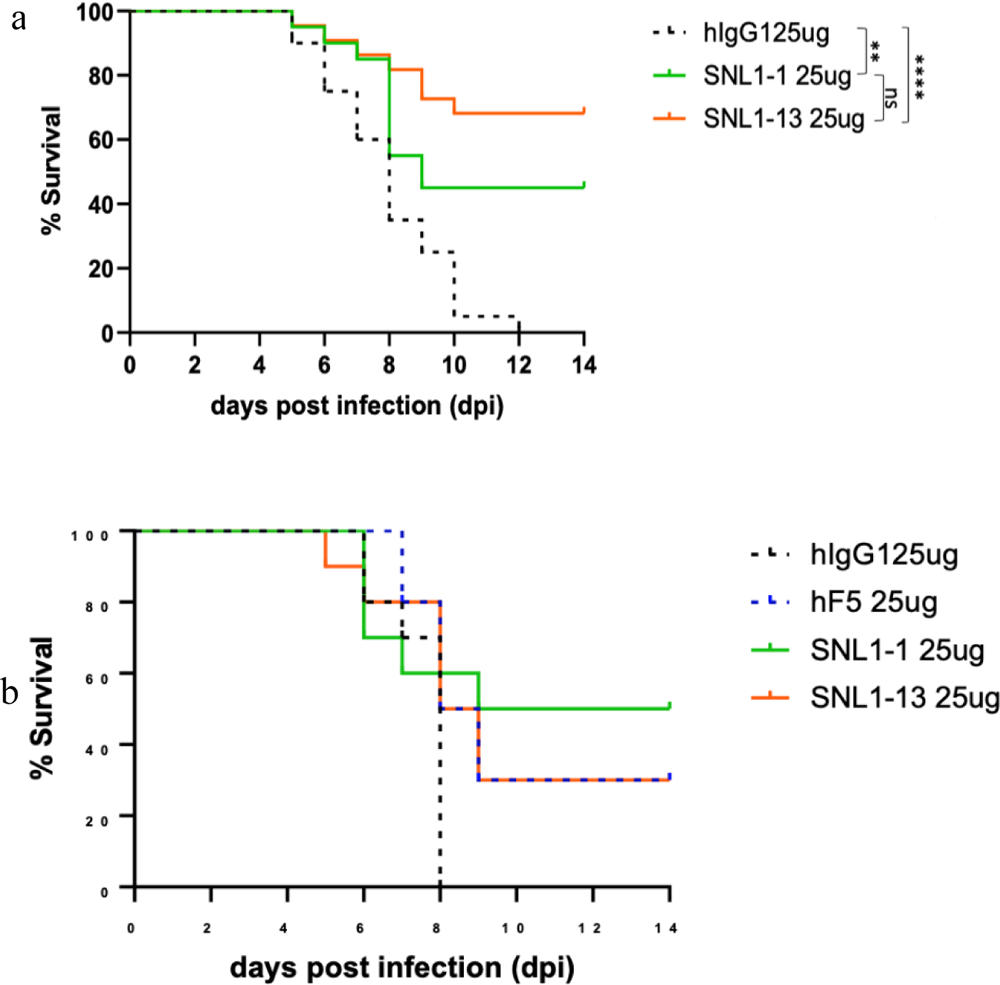
Parental and and modified antibodies antibodies display similar therapeutic efficacy during lethal VEEV TC-83 infection. Mice (n=10) were infected intranasally with 5x10^7^ PFU VEEV TC-83 and dosed with hF5, SNL 1-1, SNL 1-13Abs1, Abs13, or isotype control antibodies at +24hr (a) and +48hr (b) post infection. Morbidity and mortality were assessed daily. **p<0.01, ****p<0.0001.

**Table 1.**
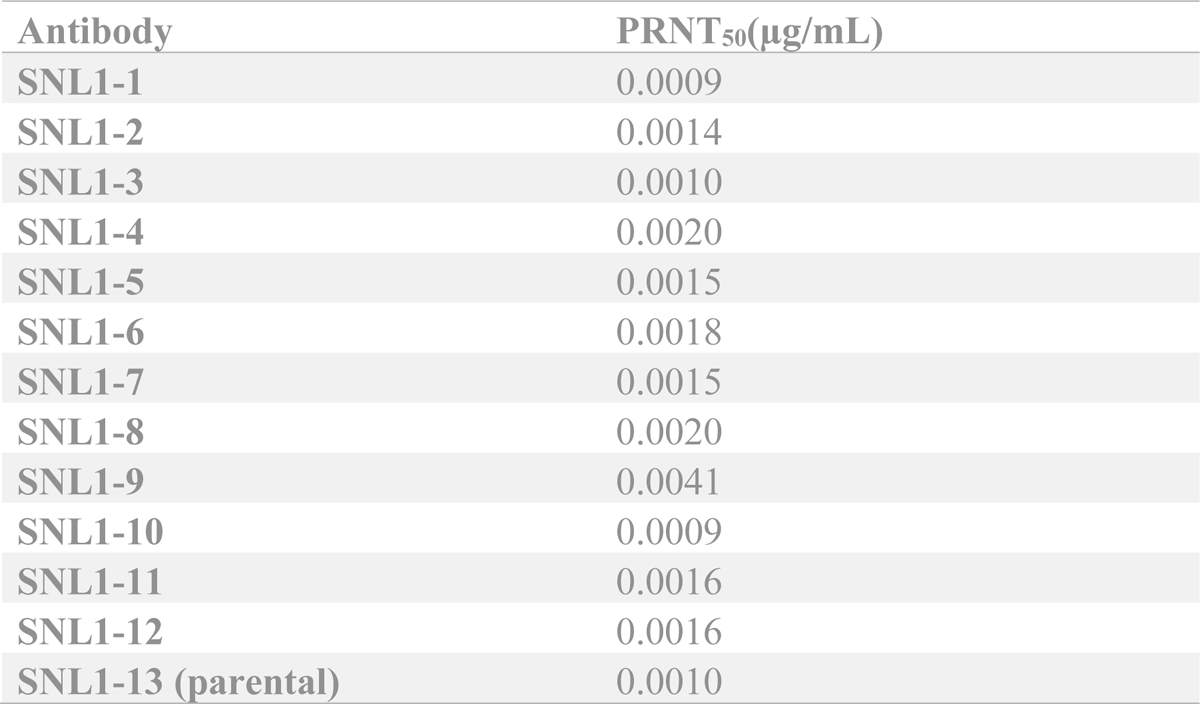
Plaque Reduction Neutralization Test data for IgGs.

## V. DISCUSSION

The combination of in-silico analysis with directed library screening led to improvement in binding to the recombinant E1E2 heterodimer by a factor of 60. For our top candidate SNL1-1, this was primarily due to a modest decrease in the Ka (association) paired with a more significant decrease in the Kd (dissociation). Library testing of nearly all possible combinations of the specified mutations was critical as it enabled determination of the value of each mutation separately as well as the value of all combinations. Combining all predicted mutations into a single construct did not result in a successful clone. For the clones that had improved binding to the target, the mean number of mutations was 3.8. We expect that the predictive power will improve with better structural data and more accurate modeling. The directed library clearly outperformed the random library, identifying nine advantageous mutations versus two advantageous mutations for the random library. One of the beneficial mutations in the random library was in the framework region which was not addressed in the directed library. Sequence Tolerance and dTERMen contributed roughly equally in terms of successful predictions. FlexddG, employed subsequent to the experimental screening, identified all the beneficial mutations as favorable. However, only four of the experimentally-determined favorable mutations were predicted to be highly favorable by FlexddG. Had FlexddG been performed prior to library generation, most likely only those four would have been selected for inclusion in the library. We conclude that for this case Sequence Tolerance and FlexddG were of comparable value for predicting favorable mutations. Sequence Tolerance is substantially less computationally intensive and is available on the ROSIE public server.

The successful prediction of mutations to improve binding despite the absence of high-resolution structural data for the H3 loop may be partly explained by the fact that the large H3 loop likely inserts into the cavity in the center of the E1E2 trimer. In that case the computational modeling may simply have needed to successfully identify H3 residues likely to be in contact with the antigen.

While improved binding to E1E2 heterodimer was achieved through the combination of in-silico analysis and library screening, we were not able to detect improvements in the binding to TC-83 virus or in neutralization efficacy against TC-83. Given limitations in our approach which did not allow us to evaluate binding to TC-83 virus by BLI, we cannot currently determine if the mutations improved binding to E1E2 trimer in the context of virus. It is possible that that differences may have existed between the antigenic characteristics of the recombinant E1E2 heterodimer and the E1E2 trimer in the virus. This could be related to the previously reported observation that difference maps between the antibody-bound and native virus suggest that the E2 A domains contract inward by ∼ 5 Å upon binding of the F5 Fab [24]. This movement could be more (or less) constrained for the E1E2 trimer bound to the virus compared with free soluble E1E2 heterodimer and may account for the differences in binding of the IgGs for the two antigens. Alternatively, it is equally likely that the approaches we utilized to evaluate virus binding were not powerful enough to detect the differences observed in the antigen binding assays.

Ultimately, the mutations that were made to SNL1-1 which showed significant improvement in binding to purified TC-83 E1E2 antigen did not improve antibody functionality either by neutralization potency (Figure 9), or therapeutic dose after lethal challenge (Figure 10). Although we did not perform an extensive analysis of therapeutic dose or administration window, this likely suggests that the native binding activity of the parental F5 antibody is sufficient for potent neutralization and protection, and the further increases in antigenic binding has diminishing returns for therapeutic potential. We propose that a similar process could be used to improve the therapeutic effect of an antibody with a less favorable antibody-antigen interaction such as when re-targeting an existing antibody against a new emerging viral variant.

## VI. CONCLUSIONS

Through a combination of computational modeling and experimental library screening we substantially improved the affinity of F5 for recombinant E1E2 of VEEV (TC-83). This is especially significant because high-resolution structural data were not available for either F5 or VEEV and the H3 loop of F5 was comprised of 20 residues which posed a severe challenge for structural modeling. However, while the approach led to improvement in binding to recombinant E1E2, the improvement did not translate to improved binding to the virus or to in-vivo efficacy against VEEV. Although this approach did not generate more potent protective antibody variants in this case, it may be useful in the future for improving low affinity Abs against known targets or for re-targeting existing therapeutic antibodies against new emerging pathogenic variants. .

## ACKNOWLEDGMENTS

This work was supported by the Defense Threat Reduction Agency [contract HDTRA140027, project CB10489, PI: Harmon]. Sandia National Laboratories is a multimission laboratory managed and operated by National Technology & Engineering Solutions of Sandia, LLC, a wholly owned subsidiary of Honeywell International Inc., for the U.S. Department of Energy’s National Nuclear Security Administration under contract DE-NA0003525. All work performed at Lawrence Livermore National Laboratory is performed under the auspices of the U.S. Department of Energy under Contract DE-AC52-07NA27344. This paper describes objective technical results and analysis. Any subjective views or opinions that might be expressed in the paper do not necessarily represent the views of the U.S. Department of Energy or the United States Government.

## Supporting information

**Figure S1.**
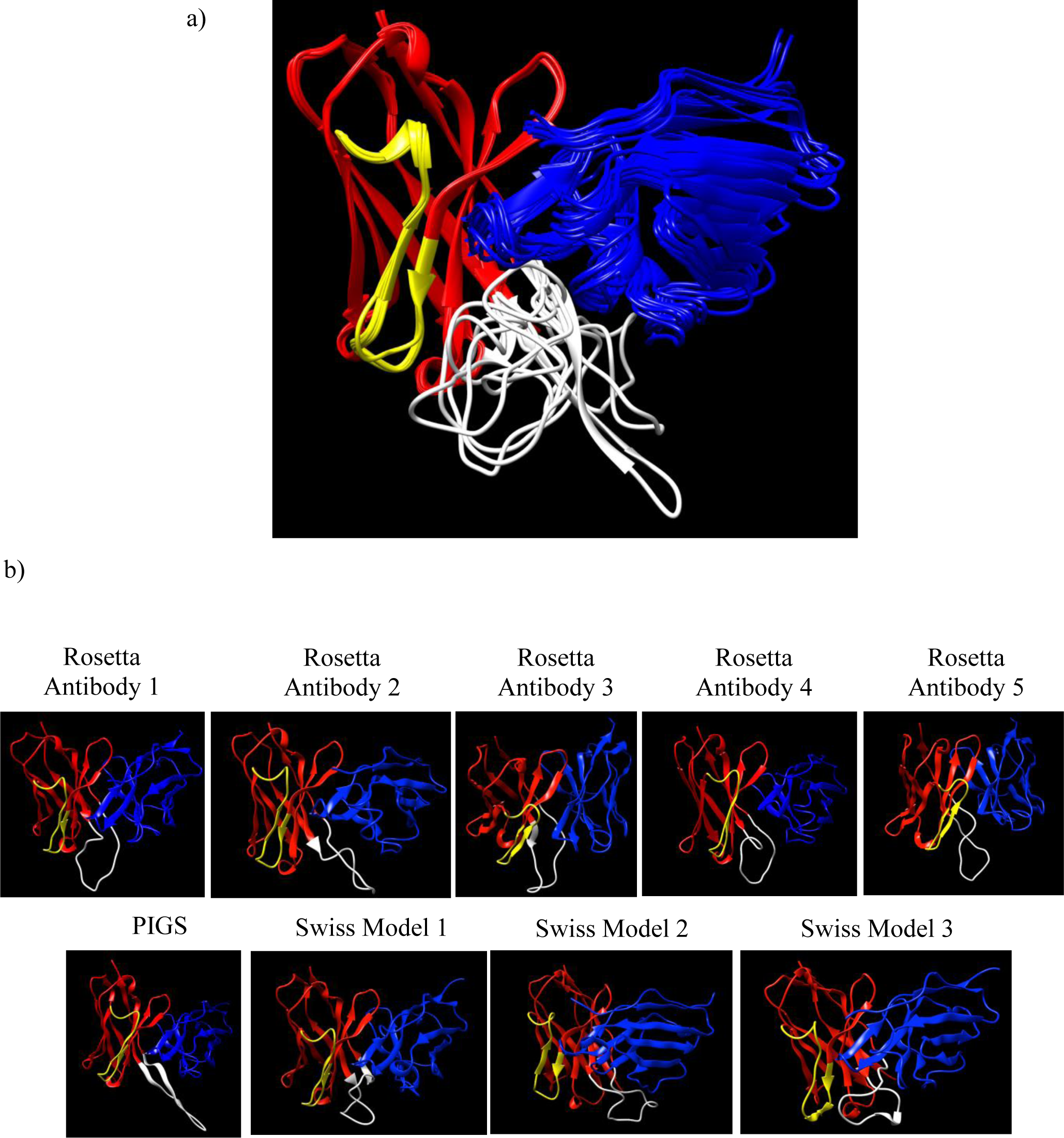
Structures of the nine F5 models used in this work, shown overlapped in a) and displayed separately in b). The H2 loop of F5 (VISHDGSHEEYADSG) is shown in yellow and the H3 loop of F5 (DGAYYYDYSGYPYDYNGIDV) loop is shown in white. The H2 loop structure is nearly identical in eight of the nine structures, but the H3 loop varies greatly among the 9 models.

**Figure s2.**
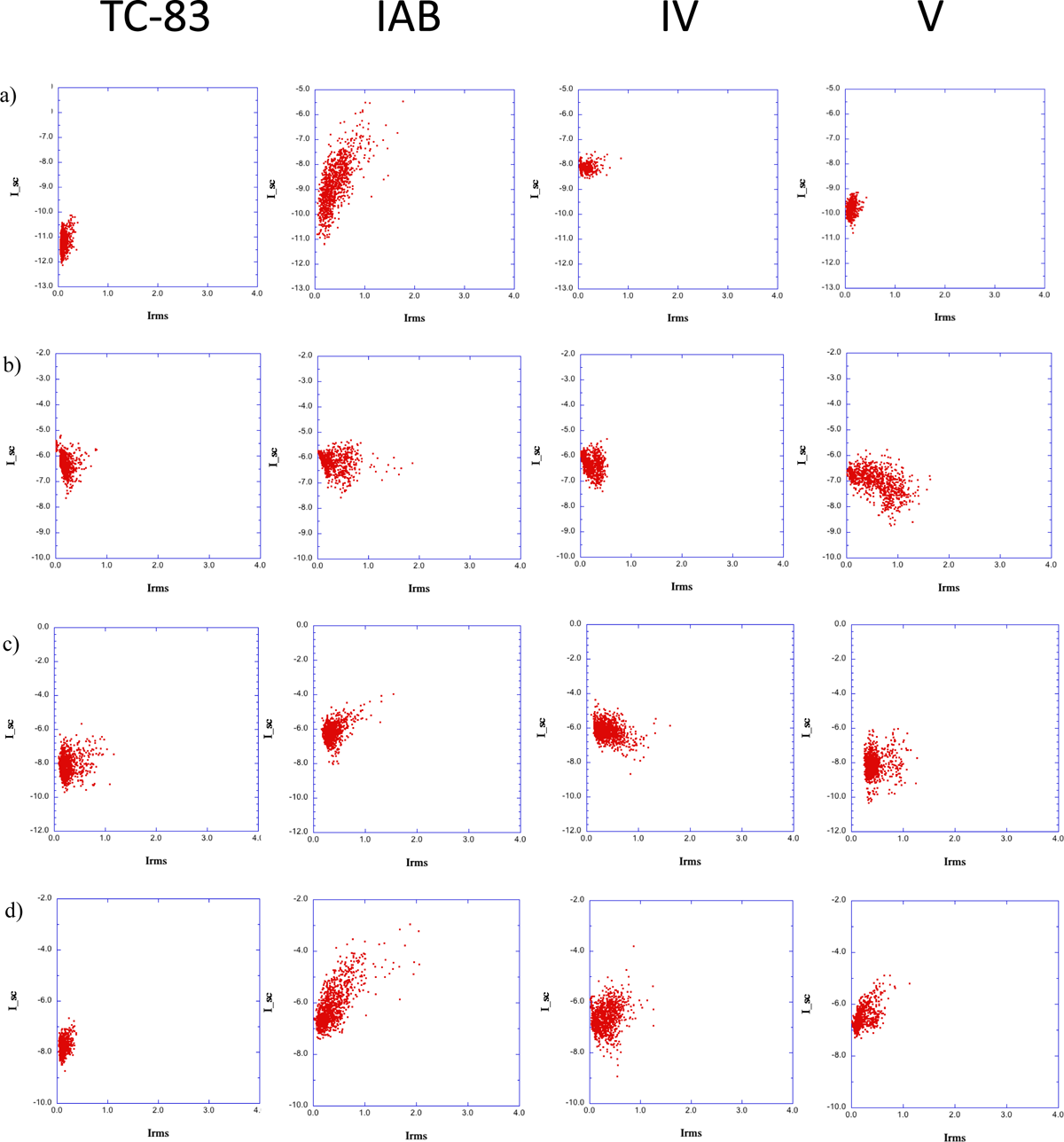
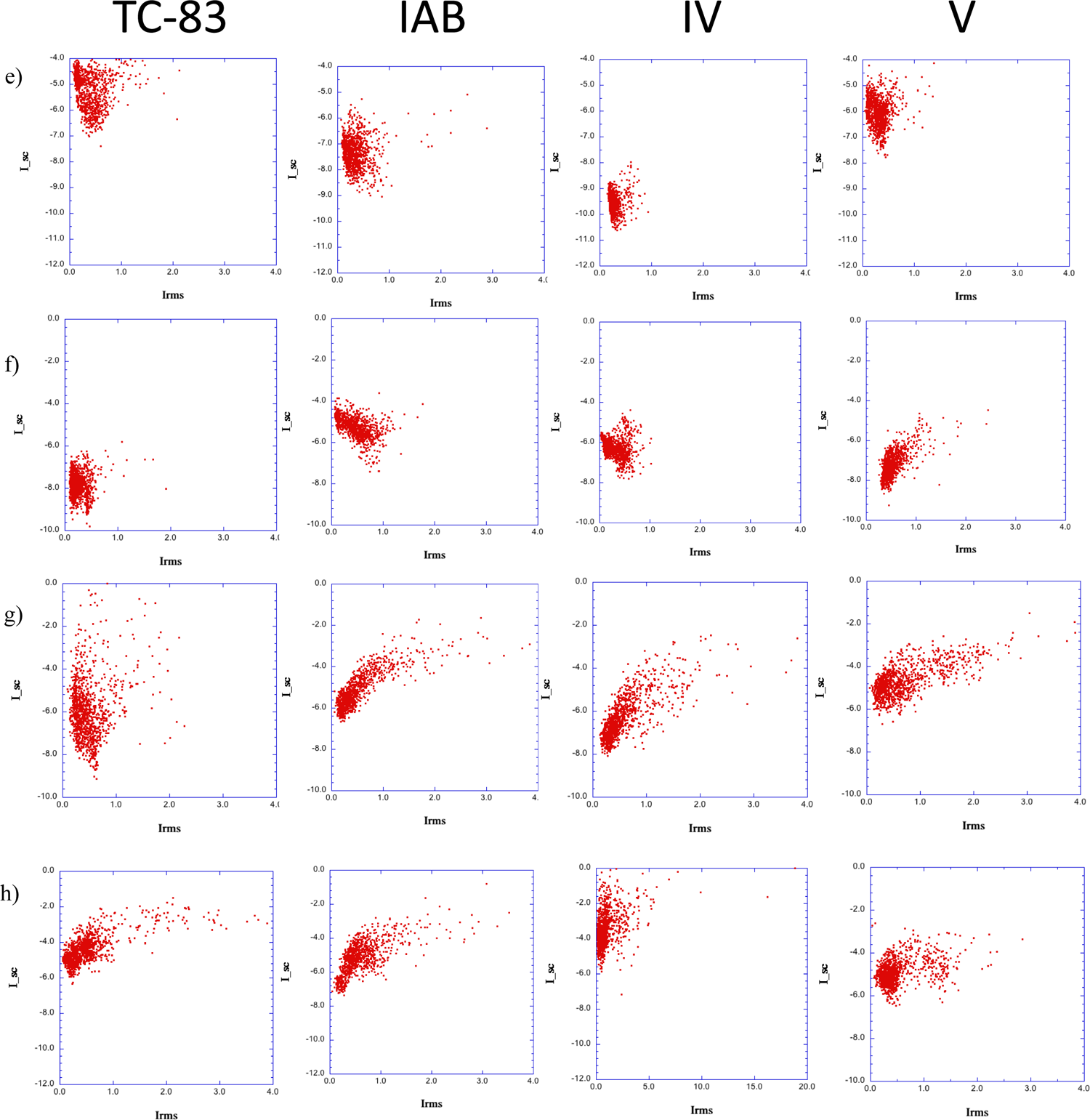
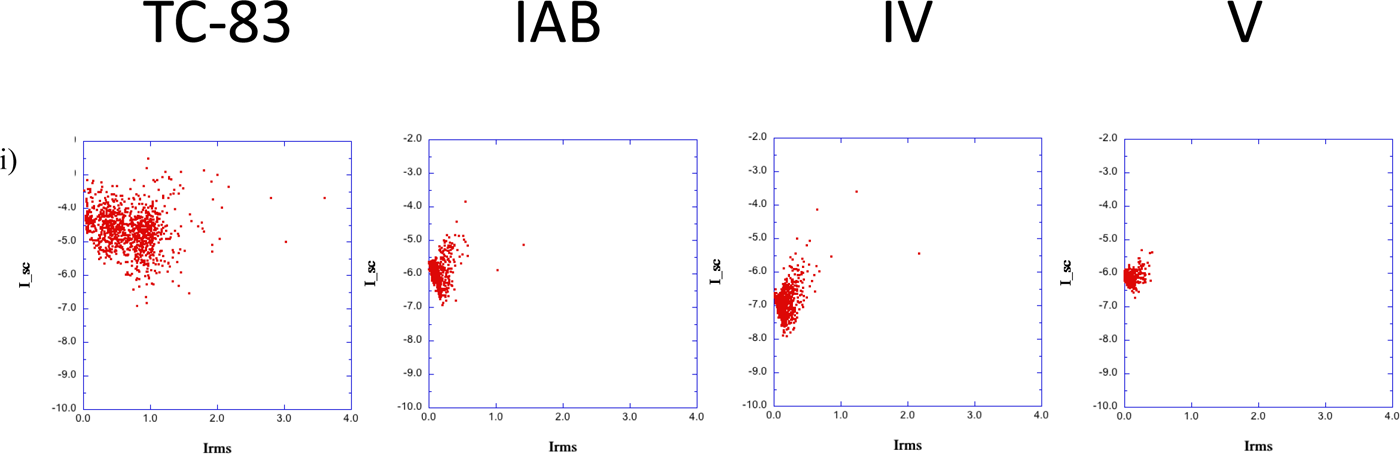
Docking scores for the following antibody structures: a) RosettaAntibody1, b) RosettaAntibody2, c) RosettaAntibody3, d) RosettaAntibody4, e) RosettaAntibody5, f) PIGS, g) SwissModel1, h) SwissModel2, i) SwissModel3. I_sc is a docking score for the interface region where lower values indicate stronger binding and lower energy. Irms is the root mean square deviation of Cα atoms from the initial configuration.

**Figure S3.**
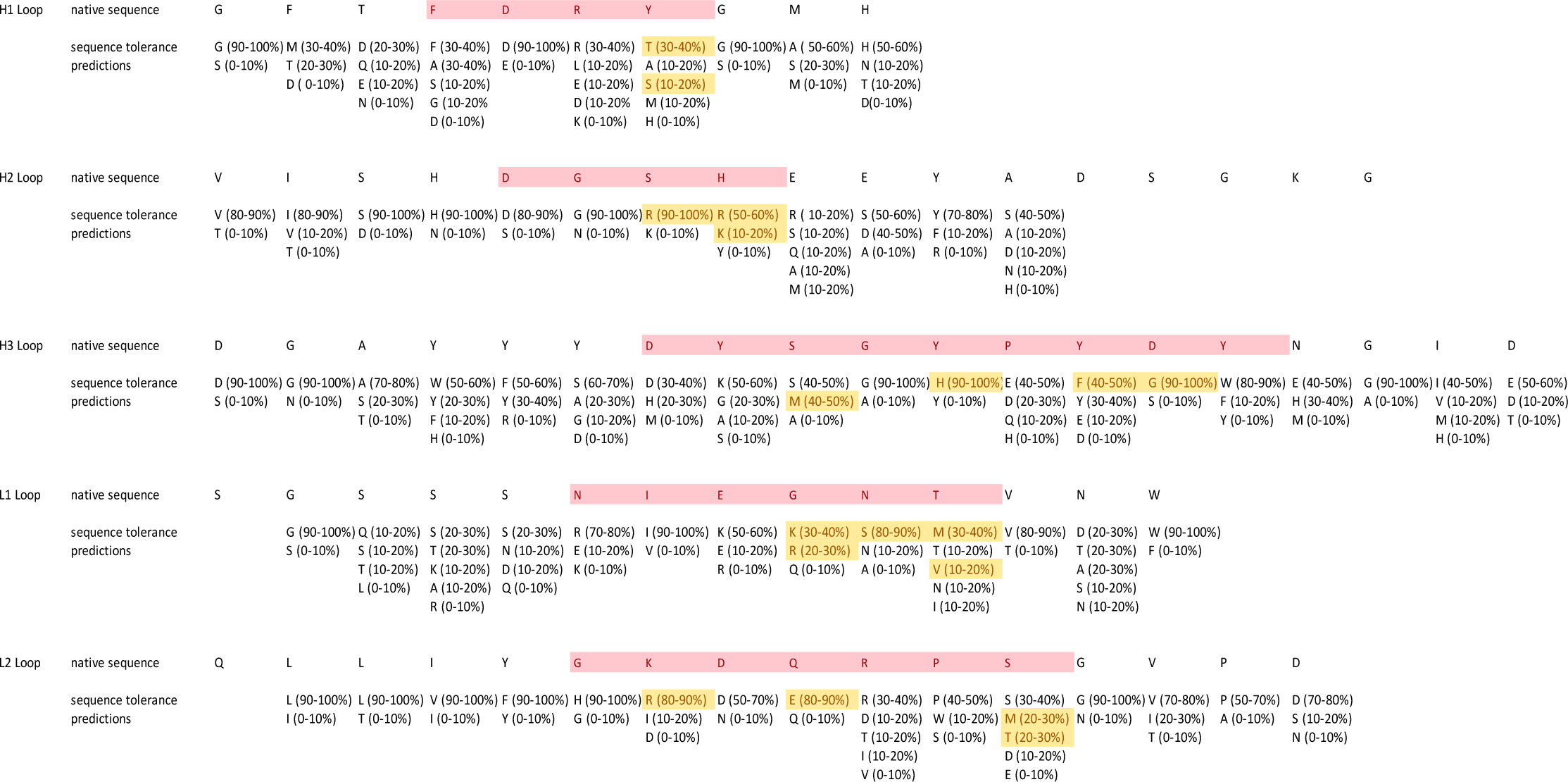
Results of Sequence Tolerance analysis of F5 residues in the CDR loops in the final refined F5/VEEV complex. The residues that contact the antigen are shown in pink. L3 does not contact the antigen in the modeled structure. The mutations chosen for inclusion in the experimental library are highlighted in yellow.

**Figure S4.**
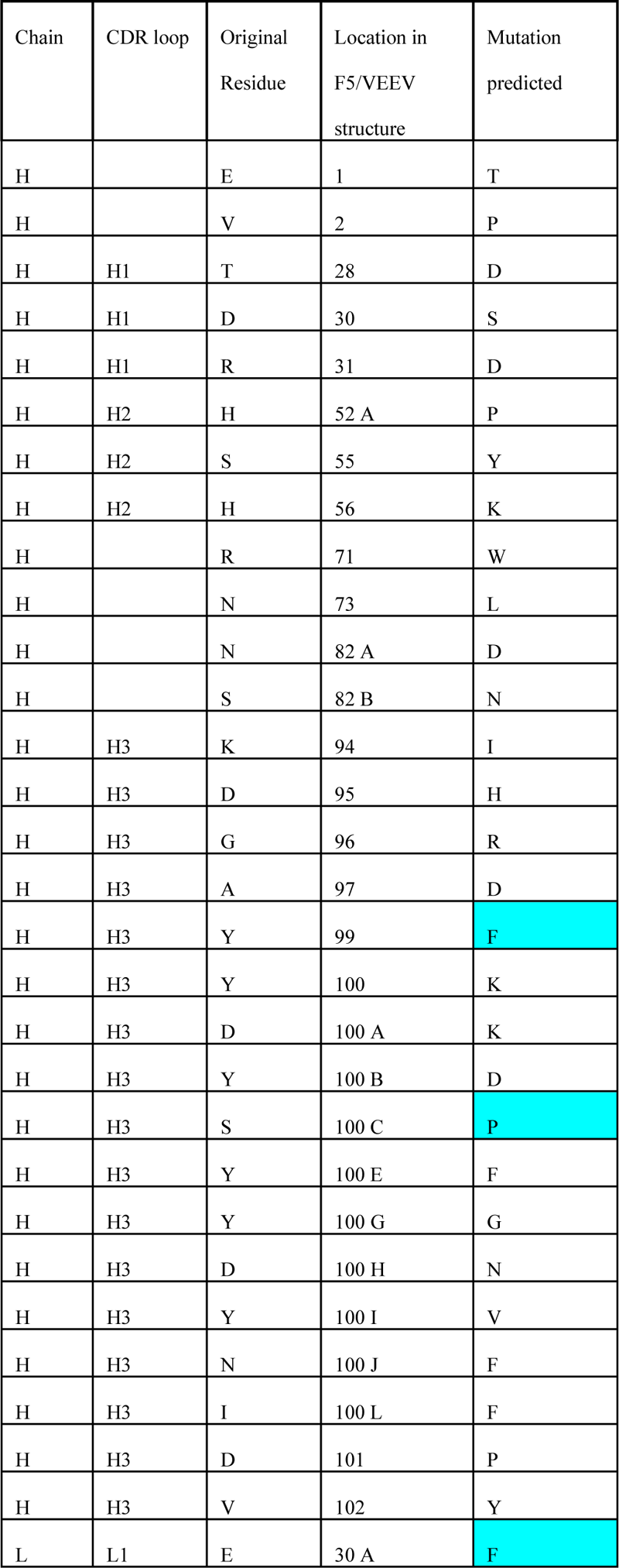
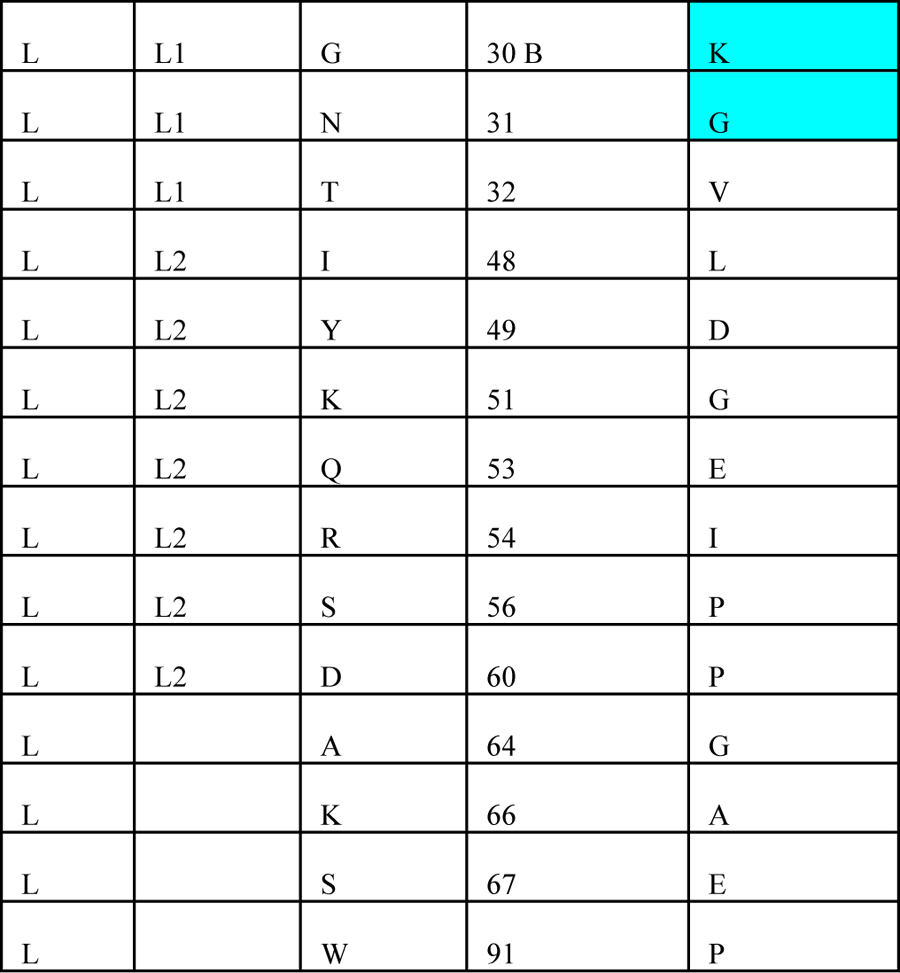
Results of dTERMen analysis of F5 residues in the CDR loops in the final refined F5/VEEV complex using pseudo-energy alone. Differences from the predictions with specificity cutoff are highlighted in cyan.

**Figure S5.**
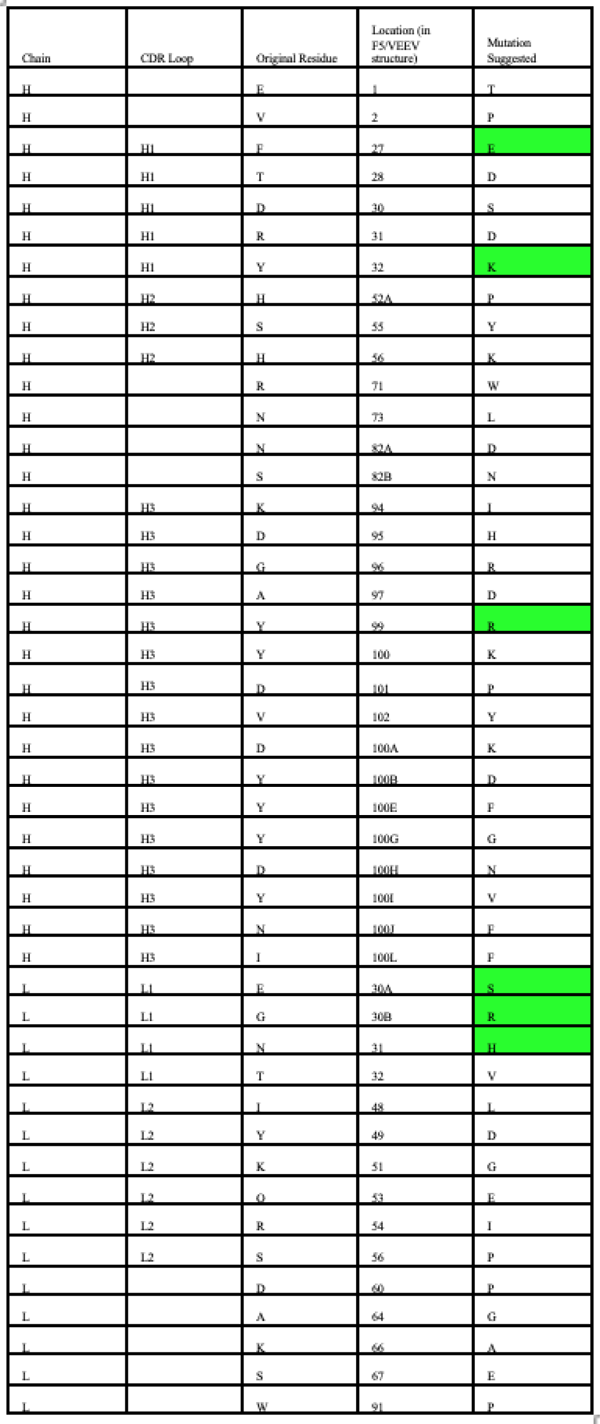
Results of dTERMen analysis of F5 residues in the CDR loops in the final refined F5/VEEV complex using energy with specificity cutoff (differences from predictions without specificity cutoff highlighted):

**Figure S6.**
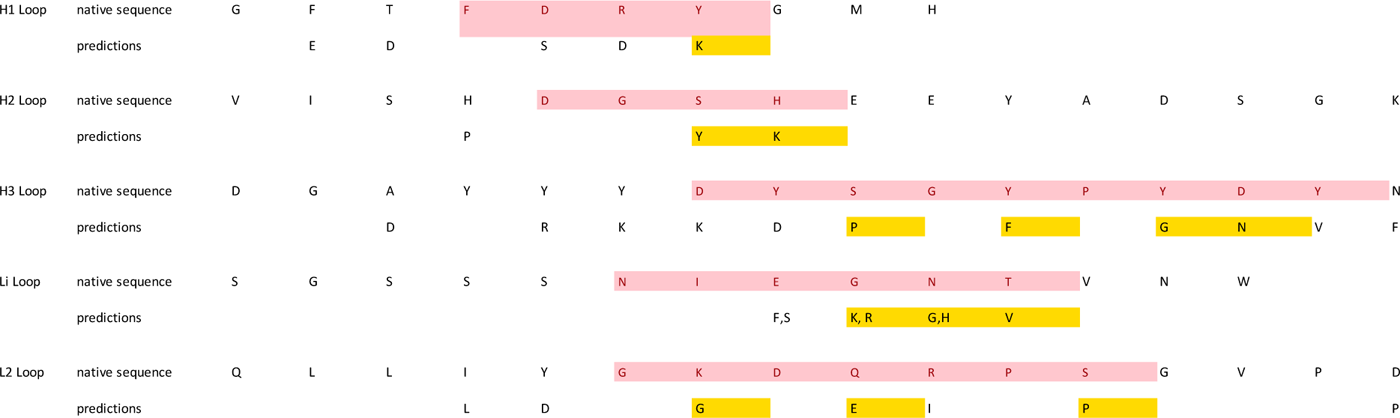
Mutations from dTERMen analysis of F5 residues in the CDR loops chosen for inclusion in the experimental library are highlighted in yellow. The residues that contact the antigen are shown in pink. L3 does not contact the antigen in the modeled structure.

**Figure S7.**
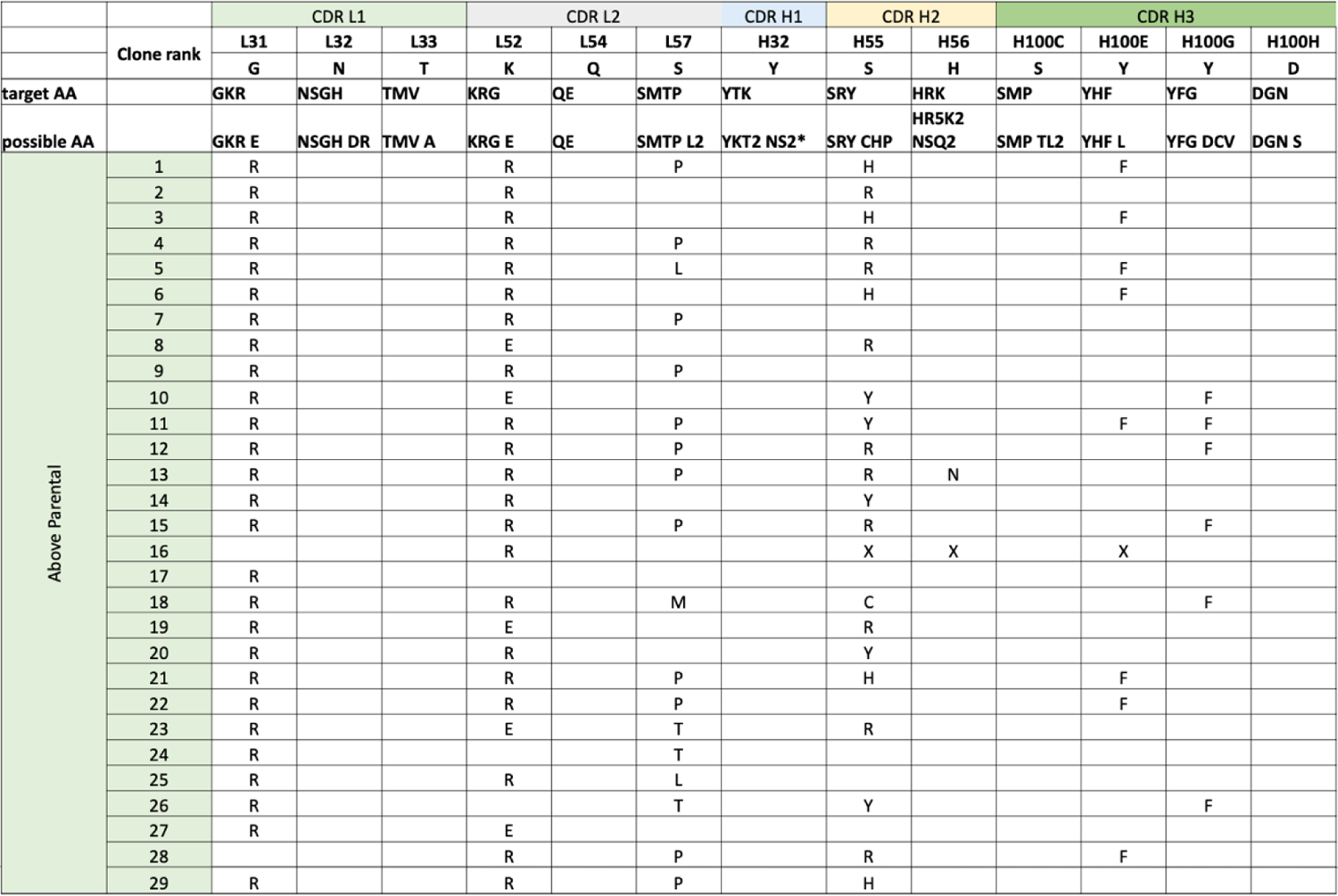
Results of experimental library screening showing mutations in clones that had ELISA signal for binding to antigen that was greater than that of the parental sequence.

**Figure S8.**
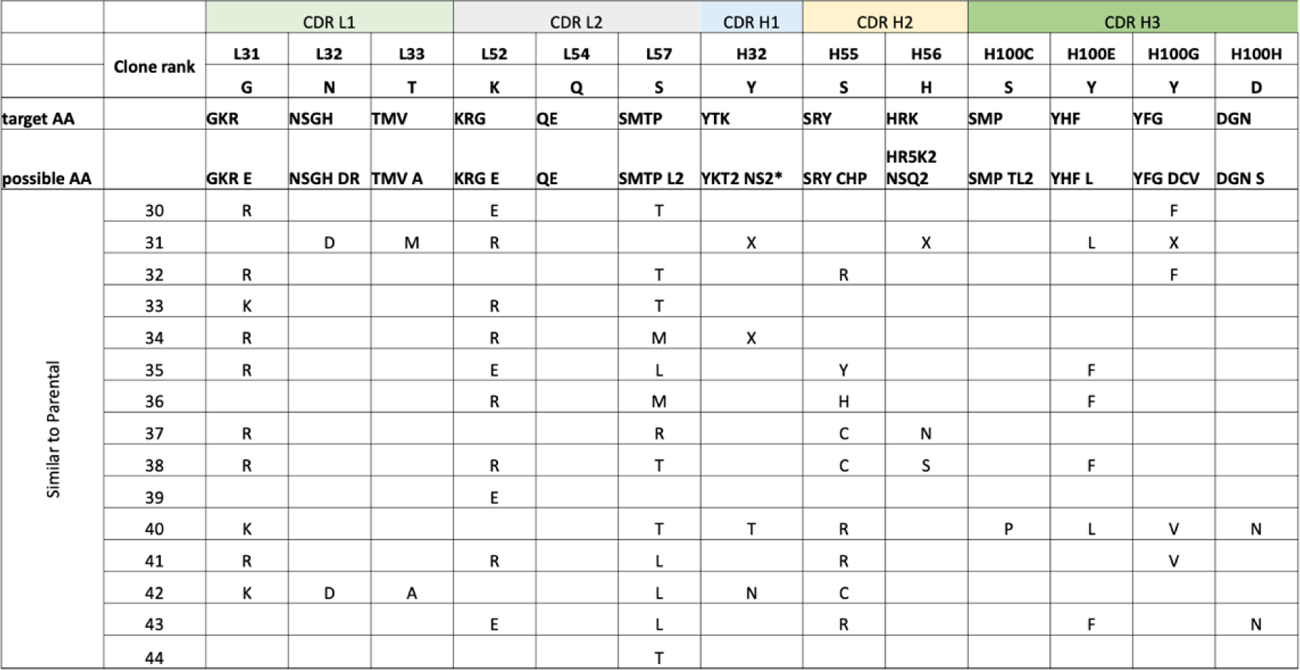
Results of experimental library screening showing mutations in clones that had ELISA signal for binding to antigen that was comparable to that of the parental sequence.

**Figure S9.**
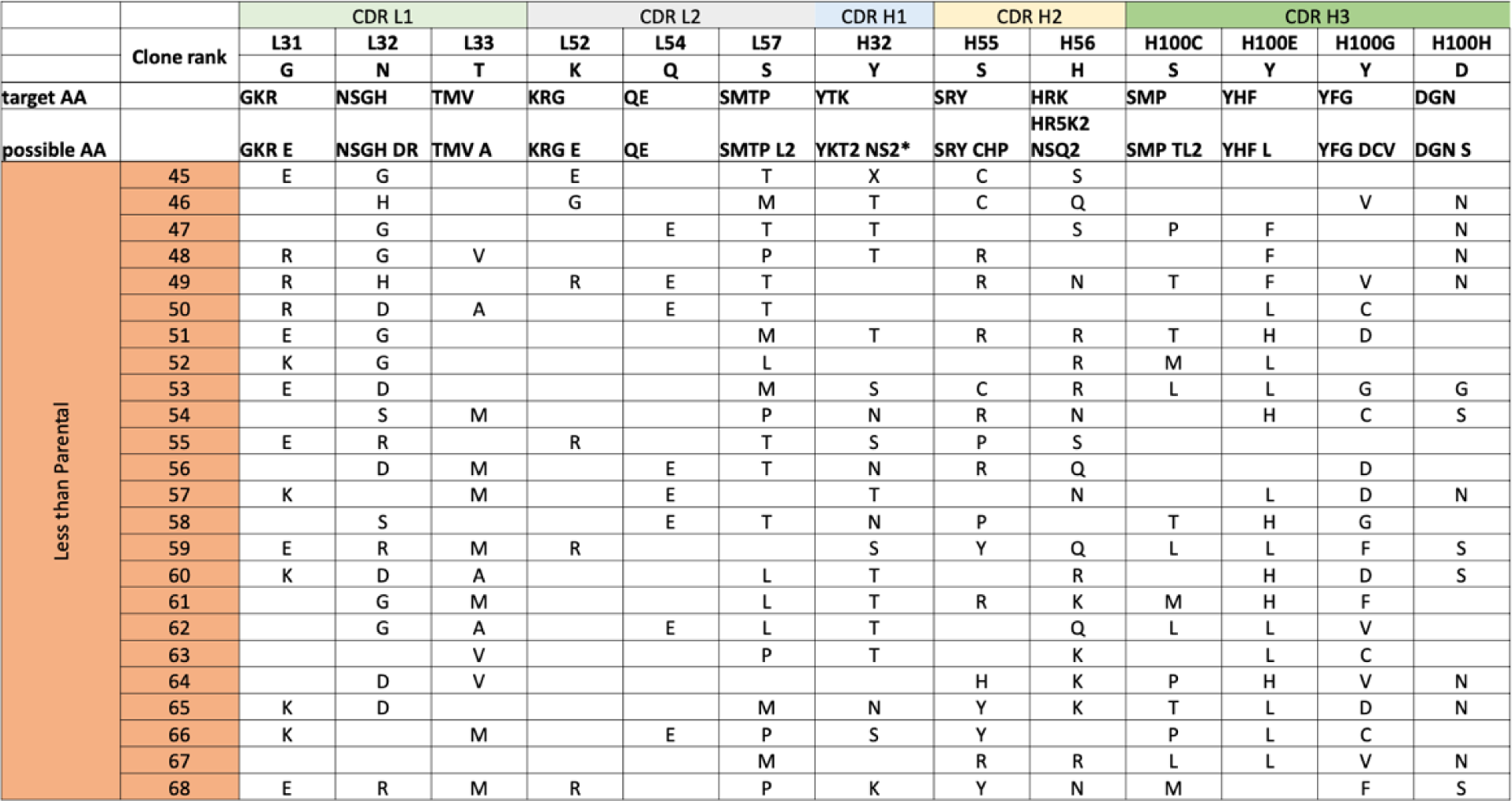
Results of experimental library screening showing mutations in clones that had ELISA signal for binding to antigen that was lower than that of the parental sequence.

**Figure S10.**
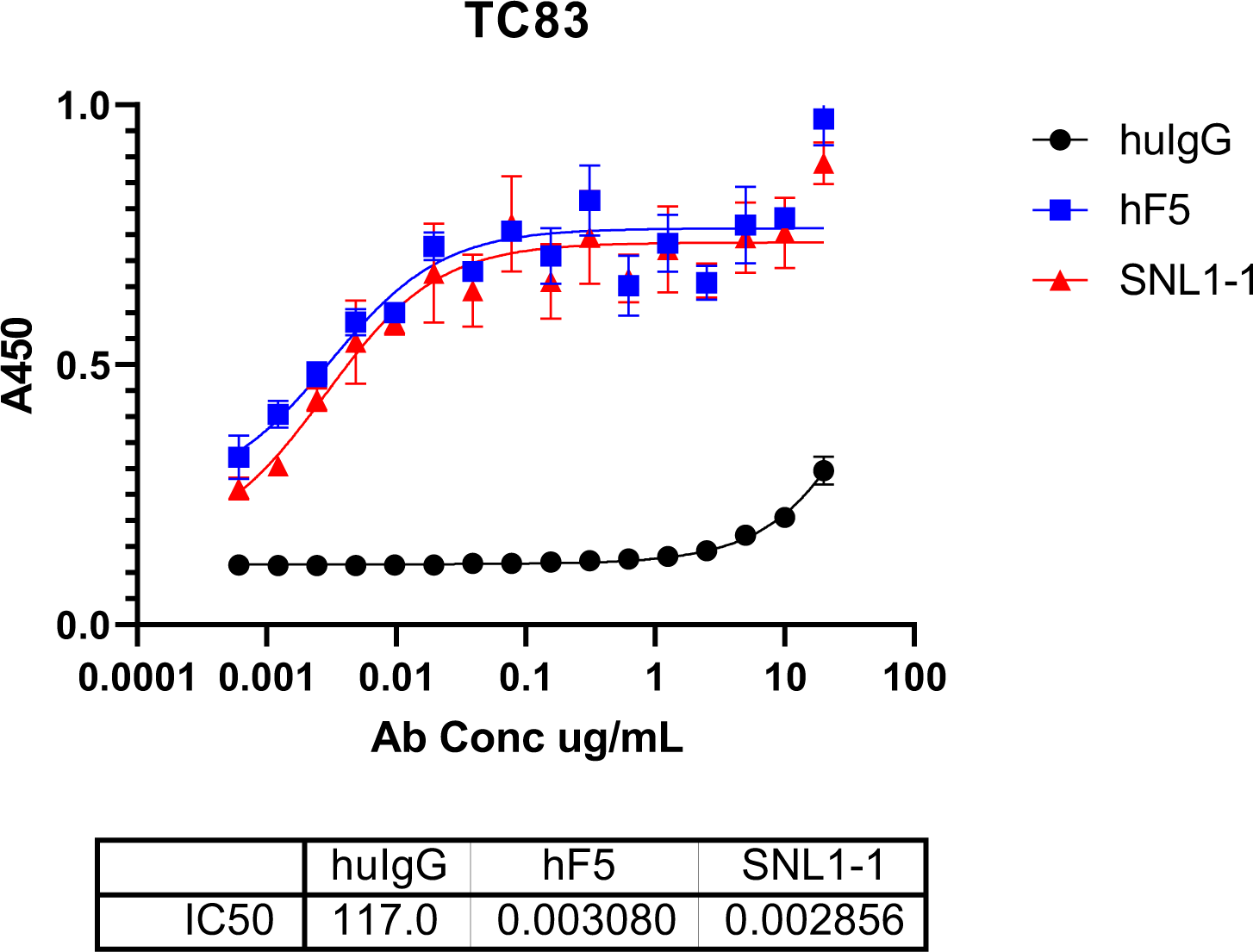
ELISA for IgGs binding to active, immobilized TC-83.

**Figure S11.**
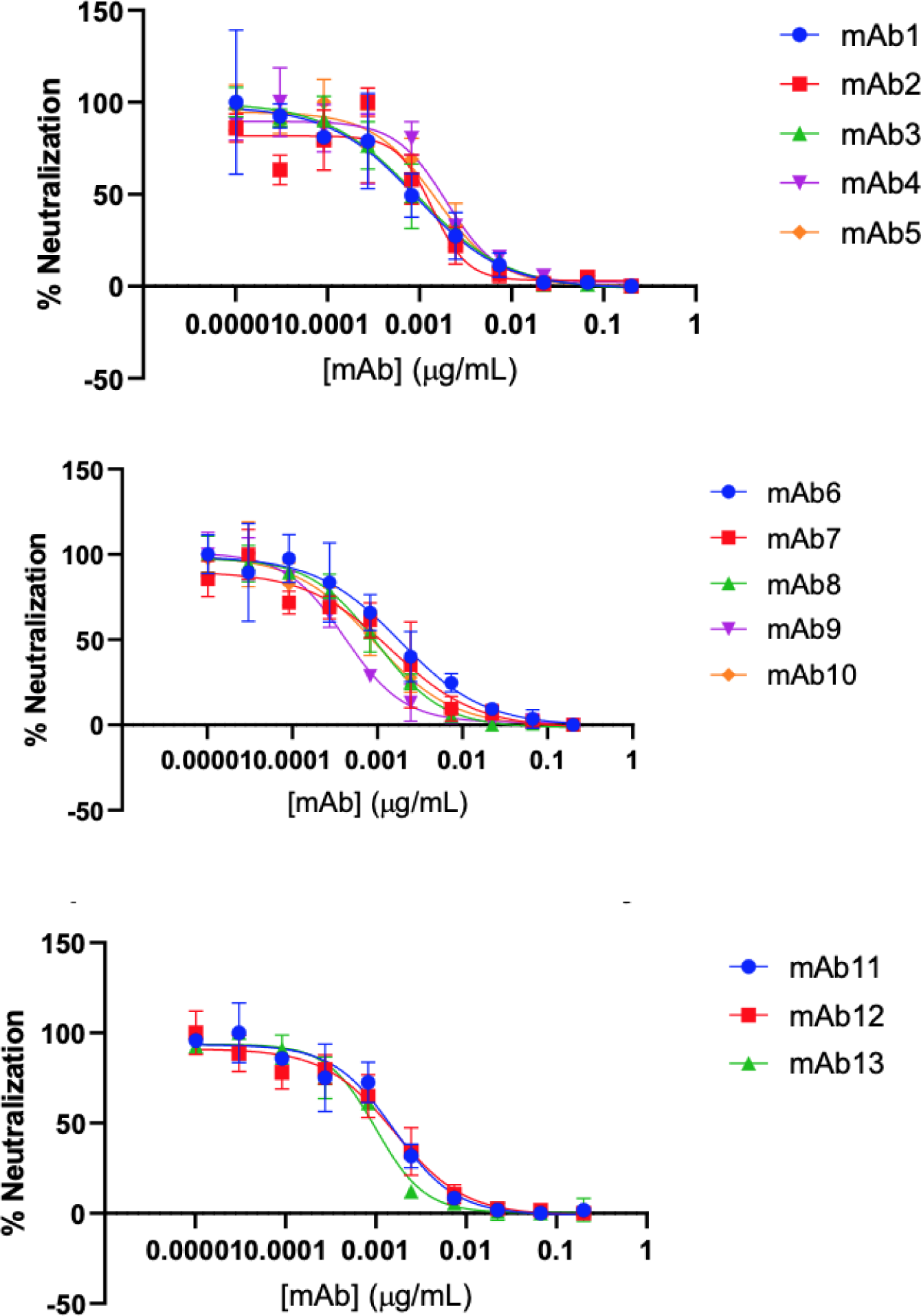
Raw data for Plaque Reduction Neutralization Test

